# Systemic 4-1BB stimulation augments extrafollicular memory B cell formation and recall responses during *Plasmodium* infection

**DOI:** 10.1101/2023.09.12.557411

**Authors:** Carolina Caloba, Alexandria J. Sturtz, Taylor A. Lyons, Lijo John, Akshaya Ramachandran, Allen M. Minns, Anthony M. Cannon, Justin P. Whalley, Tania H. Watts, Mark H. Kaplan, Scott E. Lindner, Rahul Vijay

## Abstract

T-dependent germinal center (GC) output, comprising plasma cells (PC) and memory B cells (MBC), is crucial for the clearance of *Plasmodium* infection and protection against reinfection. In this study, we examined the effect of an agonistic antibody targeting 4-1BB (CD137), a member of the Tumor Necrosis Factor Receptor Superfamily (TNFRSF), during experimental malaria. Here we show that exogenous 4-1BB stimulation, despite delaying the effector GC response, surprisingly enhanced humoral memory recall and protection from reinfection. Single cell RNA and ATAC sequencing of MBCs from mice that received 4-1BB stimulation revealed distinct populations with transcriptional and epigenetic signatures indicative of superior recall and proliferative potential. Importantly, our results indicate that the effects of 4-1BB stimulation are dependent on IL-9R signaling in B cells but independent of parasite load during primary infection. Our study proposes an immunomodulatory approach to enhance the quality of the MBC pool, providing superior protection during infection and vaccination, particularly in the context of malaria.

## Introduction

Despite decades of research, malaria remains a major public health threat, with an estimated 249 million cases and 608,000 deaths in 2022^1^. It predominantly affects poorer regions, with almost 95% of cases and 98% of deaths documented in Africa. The severity of the disease is especially prominent in individuals with limited or no prior exposure to the parasite, making children under the age of 5 the most impacted^1,2^. The newly approved anti-malarial vaccines RTS,S/AS01 and R21 provide waning efficacy even after a multi-dose regimen^3^. Clearance of the malarial parasite relies on CD4 T cell-dependent B cell activation, culminating in the production of parasite-specific antibodies during the blood stage of infection^4,5^. Prior to or concomitant with the clearance of infection, a fraction of activated B cells differentiates into memory B cells (MBCs) which quickly become antibody-secreting plasma cells (PCs) upon reinfection^6–11,12^. However, increasing evidence suggests that even after multiple exposures, naturally acquired anti-*Plasmodium* humoral immune memory affords just clinical immunity and not sterilizing immunity, resulting in repeat infections^13^.

Leveraging numerous studies in tumor models, the expression and engagement of coinhibitory receptors such as PD-1, LAG-3, and CTLA-4 on CD4 T cells have been identified as contributing to the inefficient anti-*Plasmodium* immune responses ^14–18^. In one such study, exogenous activation of OX40, a member of the tumor necrosis factor receptor superfamily (TNFRSF) expressed on T cells improved *Plasmodium* clearance following primary infection and also rendered protection from rechallenge in a model of experimental malaria ^16,17^. These studies highlight substantial room for improvement in the natural immune response against *Plasmodium* and underscore the potential of exogenous costimulatory agonists to fully harness the full breadth of the immune system.

4-1BB (CD137), another TNFRSF member predominantly expressed on T cells, has been shown to have non-redundant roles with OX40 in enhancing CD4 T cell function^19,20^. Given that CD4 T cells play a key role in anti-*Plasmodium* immunity by providing help to B cells, we investigated the role of exogenous 4-1BB stimulation in a rodent malaria model. Here, we show that exogenous ligation of 4-1BB using a monoclonal antibody significantly delayed the effector humoral response while paradoxically enhancing humoral immune memory. Transcriptomic and epigenetic analysis of MBCs from treated mice suggests they have a distinct genetic signature that facilitate superior recall and strongly points to an extrafollicular origin. Additionally, this enhanced protection following 4-1BB activation depended on IFNγ-induced expression of T-BET in B cells during the effector phase and, more importantly, IL-9:IL-9R signaling on MBCs. Collectively, our study establishes a previously unidentified pathway by which MBCs can be programmed to attain superior recall potential and identify key molecules that may be targeted to enhance vaccine efficacy especially in *Plasmodium* infections.

## Materials and Methods

### Animals, infection, treatments and parasitemia

C57BL/6 mice and *Tbx21*^-/-^ mice were purchased from The Jackson Laboratory. 4-1BB*^-/-^* mice were originally obtained from Byoug S. Kwon, Korean National University^21^ and transferred to THW and RV labs for these studies. *Plasmodium yoelli* (*Py*) clone 17XNL and *P. berghei* clone ANKA (*PbA*) were obtained from the Malaria Research and Reference Reagent Resource Center (MR4; American Type Culture Collection), stocks generated and mice infected as previously described^22^. Briefly, frozen parasite stocks (200μL aliquots) were resuspended in 4mL of normal saline or PBS (1:20 dilution) and animals were infected with 1×10^6^ parasitized RBCs intravenously. Treatments with all biologics were done intraperitoneally in a 200μL volume (in PBS) as follows: 50 μg of anti-4-1BB antibody (3H3; BioXCell), 200μg of IFNγ blocking antibody (XMG1.2; Bioxcell), 100μg IL-9 blocking antibody (9C1; Bioxcell) or rat IgG (rIgG) or mouse IgG (mIgG) on indicated timepoints. Mice were treated with artemether (Biotechne) at 20mg/kg, dissolved in 50μL of mineral oil at indicated timepoints. Parasitemia was measured by flow cytometry as previously described^23,24^. All experiments and procedures were approved by the Rosalind Franklin University of Medicine and Science Institutional Animal Care and Use Committee.

### Competitive mixed bone marrow chimeras

C57BL/6 mice (CD45.1/.1) were lethally irradiated (950 rads) and transferred with 10×10^6^ cells containing a mixture of μMT cells (CD45.1/.2) along with *Tbx21*^-/-^ (CD45.2/.2) or *Il9r*^-/-^ (CD45.2/.2) or WT (CD45.2/.2) cells at an 8:2 ratio, respectively. For CD4-specific competitive mixed bone marrow chimeras, TCRα^-/-^ were used as recipients and transferred with 10×10^6^ cells of CD4^-/-^ cells (CD45.2/CD45.2) along with *4-*1BB^-/-^ (CD45.1/.1) or WT (CD45.1/.1) cells at an 8:2 ratio, respectively. At 6 weeks post transfer mice were checked for reconstitution (>90% donor; 80% μMT or TCRα cells and 10% KO cells) by flow cytometry and at 8 weeks post transfer, reconstituted mice were used for the experiments listed elsewhere.

### Confocal imaging

Spleens were processed as previously described^22^. Antibodies used for staining are B220-AF488 (clone RA3-6B2; eBioscience), CD4-AF594 (clone GK1.5; BioLegend) and GL7-AF647 (BD Pharmingen). Imaging was done using a Zeiss LSM710 confocal microscope and images were processed using IMARIS x64 software (version 9.2.1).

### ELISpot and ELISA

For ELISPOT, Nunc MaxiSorp white polystyrene plates were coated with 0.5 μg ml^−1^ of recombinant MSP1_19_ and blocked for at least 2 h with supplemented Roswell Park Memorial Institute (RPMI) 1640 media. Bone marrow cells from *PbA*-infected mice were serially diluted in supplemented RPMI 1640 media and plated for 20 h at 37 °C with 5% CO_2_. Plates were washed with PBS + 0.05% Tween 20 and incubated overnight with horseradish peroxidase (HRP)-conjugated IgM at 4 °C. Spots were developed with 3-amino-9-ethylcarbazole.

For ELISA, Nunc MaxiSorp plates were coated with 0.5 μg ml^−1^ MSP1_19_ and blocked with 2.5% w/v BSA + 5% v/v fetal bovine serum. Serum samples from naive and *Py*-infected mice were serially diluted and incubated for 18 h at 4 °C. MSP1_19_-specific antibodies were detected with HRP-conjugated IgM, IgG2b or IgG2c at 1:1000 dilution. Plates were developed using a SureBlue reserve TMB Kit (KPL), according to the manufacturer’s protocol, and absorbance was measured at an optical density of 450 nm using a Accuris ELISA plate reader. End-point titers were extrapolated from a sigmoidal 4PL (where x is the log concentration) standard curve for each sample. The threshold for end-point titers was the mean plus 0.5–8 multiplied by the standard deviation recorded for naive mouse sera.

### Ex vivo B cell activation

Splenic cells were enriched for B cells using the MojoSort™ Mouse Pan B Cell Isolation Kit (Biolegend) according to the manufacturer’s protocol. 7.5 x10^5^ B cells were plated in triplicates and were activated with anti-IgM (10μg/mL) and anti-CD40 (10μg/mL). Some wells received rIFNγ (20ng/mL).

### Intracellular cytokine stimulation

Single cell suspension of splenocytes were prepared and 1x 10^6^ were seeded in U-bottom wells in 96-well plate. The cells were stimulated with cell activation cocktail (BioLegend) containing phorbol myristate acetate (PMA) and Ionomycin in the presence of brefeldin A (BioLegend) for 4 hours at 37°C and 5% CO_2._ After incubation, cells were stained as described elsewhere in the manuscript.

### SpyCage Reagent for detecting and enriching MSP1_-19_ specific B cells

SpyCage reagents bearing 60 copies of either mScarlet (“RedCage”) or mNeonGreen (“GreenCage”) were covalently loaded with 60 copies of the immunodominant *P. yoelii* antigen MSP-1 or the liver stage antigen *Py*UIS4 as a decoy control, respectively. Antibodies specific to SpyCage were biotinylated using amine-reactive biotinylated crosslinkers and were used to enrich SpyCage-bound cells^25^. Briefly, single cell suspensions were stained with 1.25mg of *Py*UIS4 for 10 minutes at room temperature followed by *Py*MSP-1 (1.25mg) staining for 30 minutes at 4°C. Cells were washed and incubated with 1mg of anti-aldolase biotynilated antibody for 30 minutes at 4°C. Stained cells were then washed and enriched with MojoSort™ Streptavidin nanobeads (Biolegend), according to the manufacturer’s protocol. Enriched cells were used for surface and intranuclear staining as detailed below.

### Flow cytometry

Spleens were harvested and homogenized through wire meshes to obtain single cell suspensions. After RBCs lysis, cells were filtered and counted. Samples were blocked using Fc block (clone 2.4G2) in FACS buffer (PBS + 0.002% w/v sodium azide + 2% v/v fetal bovine serum) and cells were stained with fluorescently labeled antibodies. Transcription factors were stained using True-Nuclear^TM^ Transcription Factor Buffer Set (Biolegend), according to manufacturer’s protocol. For intracellular cytokine staining Cyto-Fast Fix/Perm (Biolegend) kit was used according to manufacturer’s protocol. Samples were acquired on BD-LSR-II or Cytek Aurora and analyzed using FlowJo software (TreeStar Inc).

### MBC transfer

Memory B cells (B220^+^IgD^-^CD38^+^GL7^-^) were sort-purified from *Py*-infected mice treated with rIgG or 3H3. 2×10^5^ cells were transferred intravenously to naive μMT mice, which were infected with 1 × 10^6^ *Py*-parasitized RBCs, the following day.

### Single-cell RNA sequencing analysis

Single cell suspensions were stained with Live/Dead dye (Biolegend) and then divided in a ratio of 1:4 in which 1 part was used for sorting of polyclonal cells and 4 parts were used for MSP-1 specific cells. Before staining for surface markers, MSP-1 specific cells were stained and enriched with SpyCage reagents as previously described. Both polyclonal and MSP-1 cells were incubated with fluorescently labeled antibodies and Total-seq C antibodies (Biolegend) for 30 minutes and washed before sorting for resting B cells (B220^+^IgD^+^), polyclonal (B220^+^IgD^-^GL7^-^CD38^+^) and MSP-1 specific (B220^+^IgD^-^GL7^-^CD38^+^Decoy^-^MSP1^+^) MBCs. Post-sorted cells were run on the 10X Chromium (10X Genomics) and library preparation was performed according to the manufacturer’s protocol for Chromium Next GEM Single Cell 5’ Reagent Kit v2 with Feature Barcode technology and V(D)J amplification for mouse B cells (10X Genomics). Libraries were pooled and sequenced in the Genomics and Microbiome Core Facility at Rush University using NovaSeqX Plus (Illumina). 10X Cell Ranger Multi (v.7.1.0, 10X Genomics) was used to process gene expression, CITE-seq and V(D)J libraries in each sample individually.

Downstream analysis was performed using the Seurat package^26^ (version 5.1.0) with R (version 4.4.1). Samples were merged (rIgG and 3H3 groups in Polyclonal or MSP-1 specific pool) and the SCTransform^27^ function using the glmGamPoi^28^ package was utilized for normalization, selection of highly variable features and scaling of data. Principal component analysis (PCA) was performed and 30 dimensions were used for dimensionality reduction with Uniform Manifold Approximation and Projection (UMAP). Due to observed batch effects, the SCT datasets were integrated using the IntegrateLayers() function. Unsupervised clustering was performed using FindNeighbors() and FindClusters() function and RunUMAP() was used with reduction set to “integrated.dr”. For CITE-seq, NormalizeData() and ScaleData() were used for normalization.

Differential expression analysis was performed by PrepSCTFindMarkers() followed by FindMarkers() function, which were used to identify the markers of each cluster and differentially expressed genes (DEGs) in the 3H3 compared to rIgG group. Of the identified cluster markers, only genes with adjusted p-value < 0.05 and average fold change > 0.3 were used for the heatmaps, which show the aggregated expression of all cells within a specific cluster. Of the DEGs between 3H3 versus rIgG group, only the ones with adjusted p-value < 0.05 were kept for gene ontology (GO) analysis. Enrichment for GO analysis was performed with the clusterProfiler package (v4.12.6)^29^ with q-and p-value cutoffs of 0.05 and ontology set to “Biological Processes”. To remove redundancy of enriched terms, the function simplify() was used. To generate a gene signature score for each cell in the SCT dataset, the function AddModuleScore() was used with gene lists of GC-derived, EF-derived or atypical MBCs (**Supp. Table 1**). Single-cell trajectories (pseudotime analysis) were constructed using the Monocle3 package^30–34^. Cells were ordered by choosing the root cluster based on expression of genes associated with stemness and the earliest principal node was defined programmatically according to Monocle3’s vignette. RColorBrewer^35^, ggplot2^36^, scCustomize^37^ and viridis^38^ packages were used for visualization of the data.

### V(D)J analysis

BCR repertoire analysis was performed using Immcantation packages^39–43^. First, V, D and J genes were assigned using IgBLAST followed by removal of non-productive sequences and cells with multiple or without heavy chains. Then, cell IDs were matched with the gene expression data, cell type annotations were transferred and cells without gene expression data were removed. Clonal analysis was performed followed by generation of germline sequences. The V gene somatic hypermutation was calculated and used to compute the median mutation frequency of a clone. For visualization of data, the package ggplot2^44^ was used.

### Single-cell ATAC sequencing analysis

Polyclonal and MSP-1 specific MBCs were flow-sorted as described for scRNA-seq and nuclei was isolated. Transposition, barcoding on the 10X Chromium (10X Genomics) and library construction were performed according to the manufacturer’s protocol for Chromium Next GEM Single Cell ATAC Reagents Kits v2 (10X Genomics). Libraries were pooled and sequenced in the Genomics and Microbiome Core Facility at Rush University using NovaSeqX Plus (Illumina). 10X Cell Ranger ATAC (v.2.1.0, 10X Genomics) was used to process the data.

Downstream analysis were performed using Seurat^26^ (version 5.1.0) and Signac^45^ (version 1.14.0) packages. Cells with count lower than 500 were excluded and common peak sets (for polyclonal and MSP-1 specific MBCs) were created and filtered out based on length (10000 > peak width > 20). Peaks were quantified in each sample individually and the common set of peaks was passed as the feature argument. Samples were merged (rIgG and 3H3 groups in Polyclonal or MSP-1 specific pool) as Seurat objects and features that corresponded to sequences other than the standard chromosomes were removed. After gene annotation and based on the quality control metrics, cells were filtered based on the quantified peaks (100000 > quantified peaks > 100), frequency of reads in peaks (20%), ratio of reads mapping in blacklist regions compared to peaks (0.08), nucleosome signal (signal < 4) and transcription start site enrichment (TSS > 2). Normalization, selection of top features and dimensional reduction were performed by RunTFIDF(), FindTopFeatures() and RunSVD() functions, respectively. The first LSI component was removed since it strongly correlated to sequencing depth. Clustering was performed using the FindNeighbors() function with reduction set to “lsi” followed by FindClusters() function. GeneActivity() was used to create a gene activity matrix, which was normalized and scaled by NormalizeData() and ScaleData(), respectively. FindTransferAnchors() and TransferData() functions were used to integrate scATAC-seq data with gene expression data from the scRNA-seq. Differentially accessible regions (DARs) were calculated using the FindMarkers() function using a logistic regression (LR) model. For the total DARs, it was used a threshold of adjusted p-value < 0.05 and average log2foldchange > 0.3 and for the top DARs to be used for motif analysis it was used a threshold of adjusted p-value < 0.005 and of frequency of cells where the DAR is detected > 0.2. The list of total DARs was used to calculate the closest genes to the identified peaks, while the top DARs were used to find overrepresented motifs in the accessible regions. The list of motif position frequency matrices was obtained from JASPAR2020^46^ database. was used RColorBrewer^35^, scCustomize^37^ and ggseqlogo^47^ packages were used for visualization of the data.

### Quantitative real-time PCR

MBCs were flow sorted from *Py*-infected mice that were treated with 3H3 or rIgG and then with artemether on time points indicated elsewhere. RNA was extracted using Trizol reagent according to the manufacturer’s protocol. 2μg of RNA was used for cDNA synthesis using MMLV reagent. cDNA was added to the 2X PCR PowerUP SYBR Green Master mix at a 1:20 ratio, along with 0.2μM of forward and reverse primers. Primer sequences used in the experiments are as follows: *Gapdh* Fwd: 5’-GAG AAC TTT GGC ATT GTG G-3’; *Gapdh* Rev: 5’-ATG CAG GGA TGA TGT TCT G-3’; *Il9r* Fwd: 5’-GGA CAG TTG GCA GTA AGT CAC C-3’; *Il9r* Rev: 5’-CCA CTC TCT CCA AGG TCC AA-3’. *Ct* value were normalized to *Gapdh* using the following equation: ΔCt *= Ct (Il9r)-Ct(Gapdh).* Results are represented as a ratio of *Gapdh 2*^-(ΔCt)^

### Statistical analysis

Statistical analyses were performed using Prism 10 software (GraphPad). Specifics of each tests are detailed in the figure legends.

## Results

### Systemic 4-1BB stimulation delays effector GC response and parasite control

In our preliminary experiments with a non-lethal rodent *Plasmodium* strain, *Plasmodium yoelii* (*Py*), we observed that 4-1BB was preferentially upregulated on splenic CD4 T cells during the course of infection (**Supp Fig. 1a**). Given the upregulation of 4-1BB, we infected (WT) C57BL/6 mice or 4-1BB^-/-^ mice with *Py* and tracked the parasite load. As shown in **Supp Fig. 1b 4-1BB**^-/-^ mice had an elevated parasite load compared to WT mice, suggesting a role for this receptor in anti-*Plasmodium* immune response. We next sought to investigate the effect of exogenous ligation of 4-1BB during infection. To this end, we infected WT mice with *Py* and treated them with an agonistic monoclonal antibody to 4-1BB (3H3), or the isotype control, on 5 and 7 days post-infection (dpi) (**Fig. 1a**). We then monitored the kinetics of parasite load by measuring the frequency of parasitized RBCs during the course of infection using flow cytometry as previously described^23^. As a TNFRSF member with a co-stimulatory role like OX40^48^, we anticipated that 4-1BB stimulation would enhance the anti-*Plasmodium* humoral immune response. Contrary to this expectation, mice treated with 3H3 exhibited a prolonged infection course with an 8-fold higher total parasite load compared to controls, eventually resolving by 39-40 dpi (**Fig. 1b**). Additionally, we observed a delayed onset of germinal center (GC) responses—both GC B cells (**Supp Fig. 1c and Fig. 1c**) and GC Tfh cells (**Supp Fig. 1d and Fig. 1d**)—in 3H3-treated mice, peaking only around 39 dpi, suggesting that parasite clearance may be GC-dependent. Not surprisingly, serum parasite-specific antibody levels were also slow to accumulate in 3H3-treated mice (**Fig. 1e**). Confocal imaging of spleens from either group of mice revealed a pronounced disruption of lymphoid architecture in 3H3-treated mice, with no apparent follicular organization until 21 dpi, after which the follicles grouped into GCs as evidenced by the appearance of GL7 staining within the follicles (**Fig. 1f**). Importantly, 3H3-driven loss of parasite control was reversed in mice that lacked 4-1BB in CD4 T cells (CD4^4-1BBL-/-^) (**Supp Fig. 1e**). These data collectively demonstrate that exogenous ligation of 4-1BB on CD4 T cells in *Plasmodium*-infected mice results in derailed GC responses and heightened parasite load.

**Fig. 1:**
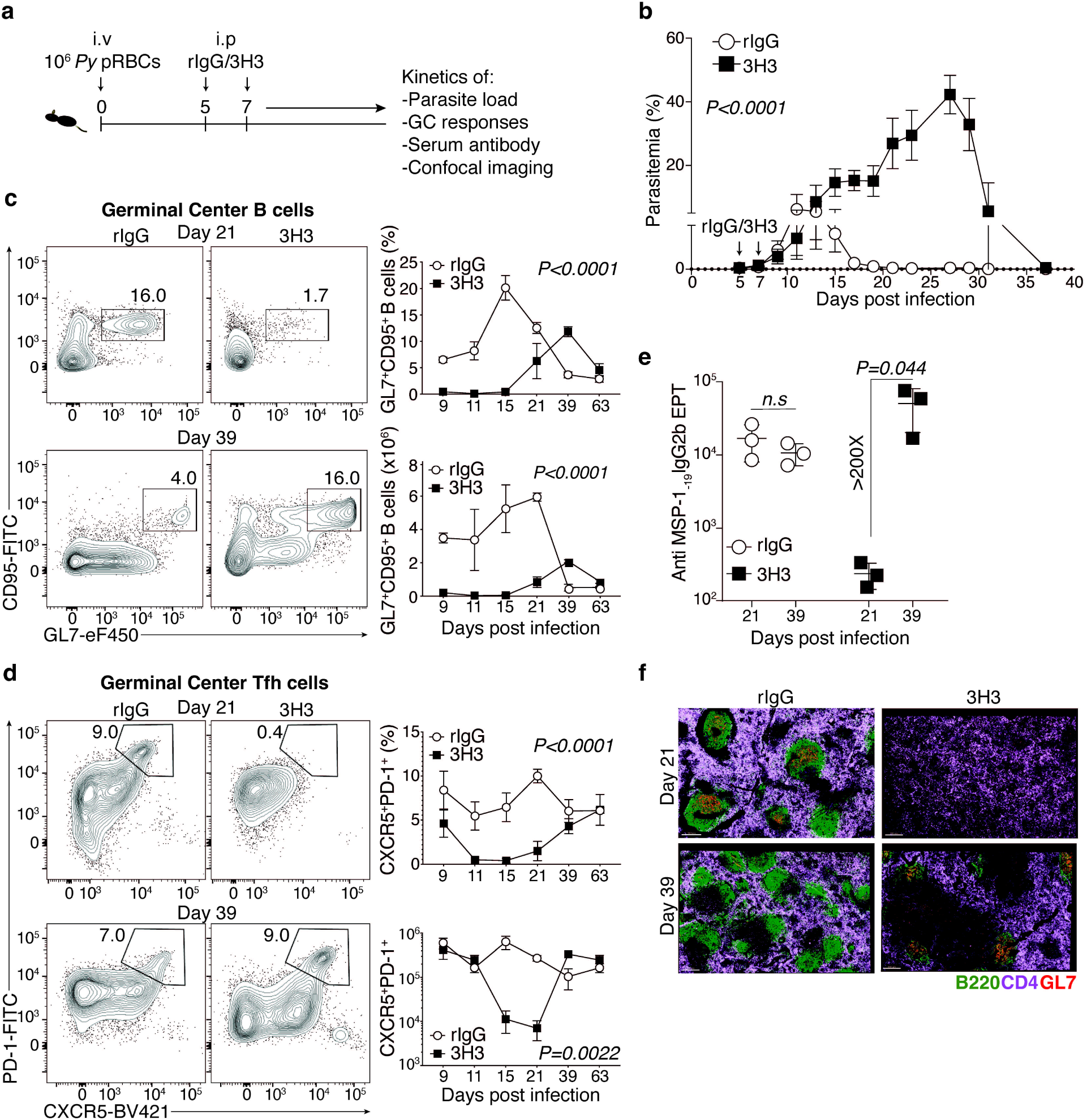
Exogenous 4-1BB stimulation leads to loss of parasite control due to derailed GC response during *Plasmodium* infection. **a,** Experimental design. C57BL/6 mice were infected with *Plasmodium yoelii*-parasitized red blood cells (*Py* pRBCs) and treated with 50μg of 4-1BB agonist antibody (3H3) or isotype control (rIgG) on days 5 and 7 post infection. **b**, Kinetics of parasite burden (% infected RBCs). Data are mean ± s.e.m and are pooled from *n=2* independent experiments using *n=7* mice for either group. **c,d,** Representative flow plots (left) and kinetics (right) of germinal center B cells (**c**) and germinal center Tfh cells (**d**) in the spleens of IgG-and 3H3-treated mice. **e,** Summary graph of anti-MSP-1_-19_ IgG2b serum antibody end-point titers (EPT) on 21 and 39 dpi. For **c-e,** data are mean ± s.d and are representative of *n=2* biologically independent with similar results using *n=3* mice for either group. **f,** Confocal micrographs of rIgG-and 3H3-treated spleens on days 21 and 39 post infection showing total B cells (green), CD4 T cells (purple) and GC B cells (red). Data are representative of at least 2 biologically independent experiments. For **f**, scale bar: 200 μm. For **b, c** and **d,** two-way ANOVA and for **e,** two-tailed Students’ *t* test was used for statistical analysis.

### Enhanced protection following rechallenge in mice exposed to systemic 4-1BB stimulation

Infection of mice with *P. berghei ANKA* (*PbA*) is invariably lethal even in mice that recovered from *Py* infection. This challenge protocol is commonly used and is considered a stringent test for MBC recall. Given that exogenous 4-1BB ligation delayed GC responses and increased the parasite burden during the primary *Py* infection, we hypothesized that mice treated with 3H3 would exhibit impaired memory responses and, consequently, reduced protection upon challenge. To test this, we monitored survival in *Py*-infected mice treated with either 3H3 or rIgG after challenge with *PbA* on day 90 post *Py* infection, when both groups were in convalescence (**Fig. 2a**). Surprisingly, 3H3-treated mice exhibited significantly longer survival compared to the rIgG group (**Fig. 2b**). To directly measure MBC recall, we assessed the fold change in antibody titers between day 0 and day 5 post *PbA* challenge. Consistent with the enhanced survival of 3H3-treated mice, we observed a greater fold increase in anti-merozoite surface protein 1 (MSP-1_-19_)-specific antibody (IgM) titers between day 0 and day 5 post *PbA* challenge (**Fig. 2c**), indicating an enhanced MBC recall response. An increase in parasite-specific antibody titers during this short window would indicate that the antibodies are derived from PCs differentiating from an existing MBC pool, rather than from a *de novo* GC reaction. The higher parasite load observed during the primary *Py* infection following 3H3 treatment (**Fig. 1b**) raises the possibility that the enhanced protection following *PbA* challenge may result from elevated inflammation and/or residual antigen load. To investigate whether these factors influenced the outcome, we truncated the primary infection and the initial parasite load by treating mice with the antimalarial drug artemether at indicated time points (**Fig. 2d**). As expected, artemether treatment resulted in complete parasite clearance in both groups, with 3H3 group showing no evidence of a productive GC response even by 21 dpi, while rIgG group retained the GC B cell and GC Tfh cells (**Supp Fig. 2a-c**) as was seen when mice were not treated with artemether (**Fig. 1**). Upon challenge with *PbA* on day 38 post-*Py* infection, we observed significantly enhanced protection in 3H3-treated mice that received chemoprophylaxis (**Fig. 2e**). Correspondingly, these mice also showed a higher fold change in anti-MSP1 antibody titers post *PbA* challenge (**Fig. 2f**). To further test whether 3H3-mediated protection is relevant to attenuated *Plasmodium* parasite vaccination, we immunized mice with irradiated blood-stage *Py* parasites and administered either 3H3 or rIgG on days 5 and 7 post-infection. On days 30-35 post-vaccination, the mice were challenged with *PbA* and monitored for survival (**Fig. 2g**). As with live parasite infections, vaccination with irradiated parasites also conferred superior protection when mice were treated with 3H3 (**Fig. 2h**), which correlated with an enhanced MBC recall response, as evidenced by higher fold changes in serum antibody titers (**Fig**. **2i**).

**Fig. 2:**
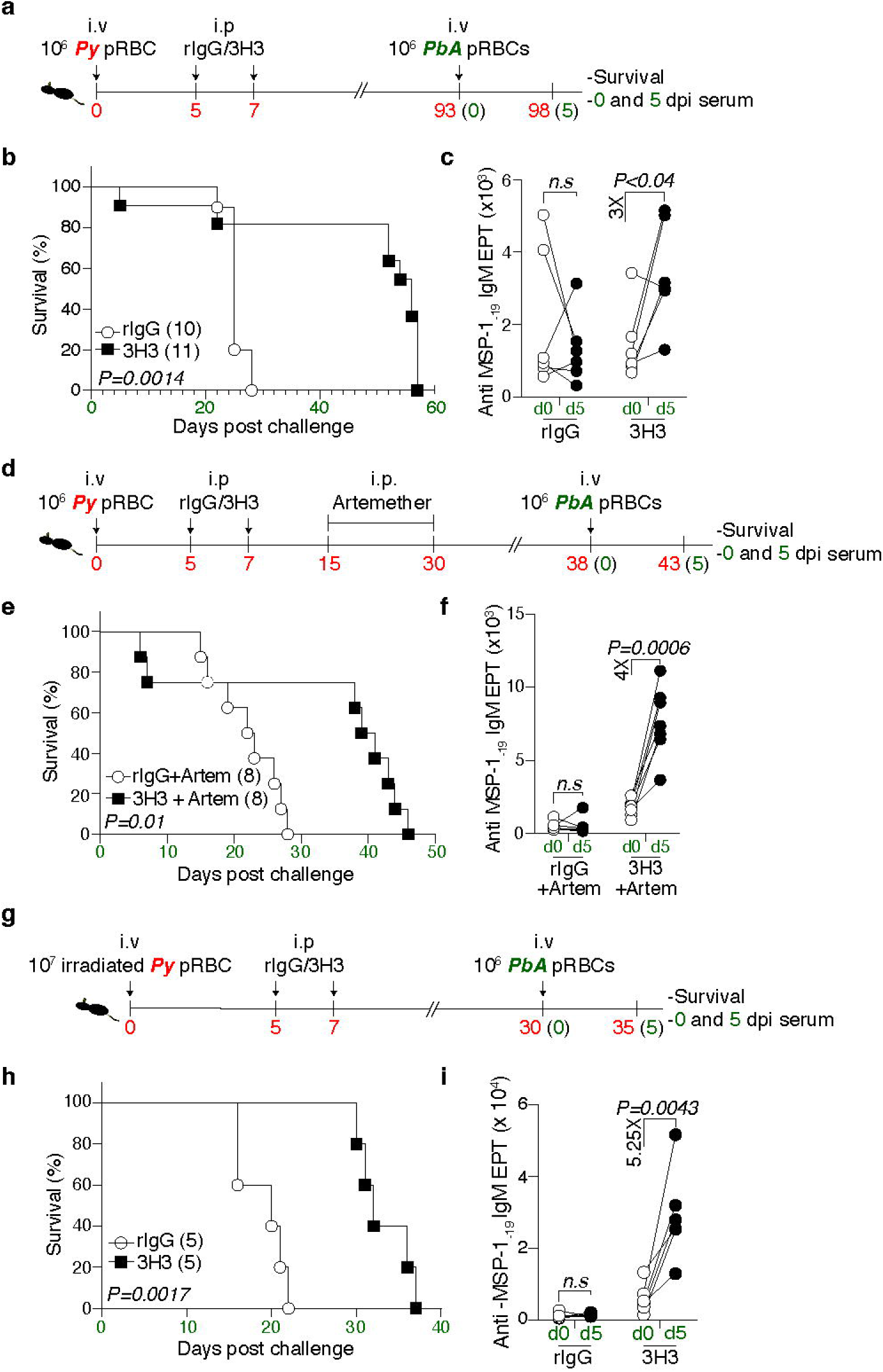
4-1BB stimulation enhances MBC recall and protection against rechallenge independently of antigen load. **a,** Experimental design. C57BL/6 mice were infected with *Py* and treated with 3H3 or isotype control as previously described. During the convalescent phase, mice were challenged with *Plasmodium berghei* ANKA-parasitized (*PbA*) RBCs. **b,** Survival graph of mice following *PbA* challenge. Data are pooled from two independent experiments. **c,** Fold change in anti-MSP-1_-19_ IgM serum antibody EPT between 0 and 5 dpi. Data are pairwise comparison of the two time points in each mouse and are pooled from *n=2* biologically independent experiments using *n=6* (rIgG) and *n=5* (3H3). **d,** Experimental design. C57BL/6 mice were infected with *Py* and treated with 3H3 or isotype control as previously described. Mice were treated with the Artemether (Artem) to control the parasite load and challenged with *PbA*. **e,** Survival of mice following *PbA* challenge. Data are pooled from two independent experiments. **f,** Fold change in anti-MSP-1_-19_ IgM serum antibody EPT between 0 and 5 dpi. Data represents pairwise comparison of the two time points in each mouse and are pooled from *n=2* biologically independent experiments with similar data using *n=7* (rIgG) and *n=7* (3H3). **g,** Experimental design. C57BL/6 mice were infected with irradiated parasites and treated with 3H3 or isotype control as previously described. During convalescence mice were rechallenged with *PbA*. **h,** Survival of mice following *PbA* infection. Data are pooled from two independent experiments. **i,** Fold change in anti-MSP-1_-19_ IgM serum antibody EPT between 0 and 5 dpi. Data are pairwise comparison of the two time points in each mouse and are pooled from *n=2* biologically independent experiments using *n=5* (rIgG) and *n=5* (3H3). For **b**, **e** and **h**, data were analyzed by Mantel-Cox test. For **c, f** and **i**, two-tailed Mann-Whitney U tests were used for statistical analysis.

In summary, these data demonstrate that systemic 4-1BB ligation, while delaying the effector GC response, significantly enhances humoral immune memory. This enhancement in memory function appears to be independent of early antigen load and holds true even when attenuated parasites are used.

### Enhanced humoral immune memory stems not from larger number but higher recall potential of MBCs

To verify whether the higher fold change in parasite-specific antibodies on day 5 post *PbA* challenge was due to larger number of antibody secreting cells (ASC), we enumerated PCs in the bone marrow of *Py*-recovered 3H3-and rIgG-treated mice challenged with *PbA.* As expected, we observed significantly larger numbers of MSP-1 specific IgM^+^ PCs in 3H3-treated mice (**Fig. 3a**). As MBCs are the precursors for PCs upon antigen re-exposure^10^, we determined the frequency and number of MBCs (IgD^-^CD38^+^GL7^-^B220^+^) prior to *PbA* challenge on 90 dpi. Contrary to our expectation, we observed significantly lower numbers of both polyclonal and antigen-specific (MSP-1 specific) (**Supp Fig. 3a, b**) MBCs in 3H3-treated mice compared to rIgG-treated ones. Upon further characterization of the MBC pool using previously described markers such as CD73, CD80 and PD-L2^49,50^ to identify functionally distinct subsets, we observed significantly lower number of polyclonal CD73^+^CD80^+^ (**Fig. 3b**), CD73^+^PD-L2^+^ and CD80^+^PD-L2^+^ (**Supp Fig. 3c, d**) MBCs. Evaluation of MSP-1 specific MBCs also showed reduced numbers of CD73^+^PD-L2^+^, CD80^+^PD-L2^+^ (**Supp Fig. 3e, f**) pools. MSP-1 specific MBCs within the CD73^+^CD80^+^ fraction was also lower in 3H3-treated mice (**Fig. 3c**). The double positive MBC fraction has been shown to readily differentiate into ASCs while the double negative pool has been shown to reseed a secondary GC response upon antigen reencounter^12, 50^. Given that we see a rapid increase in antibody titers in just 5 days of *PbA* challenge (**Fig. 2a**), it is likely that the PCs may have readily differentiated from the existing double positive MBCs.

**Fig. 3:**
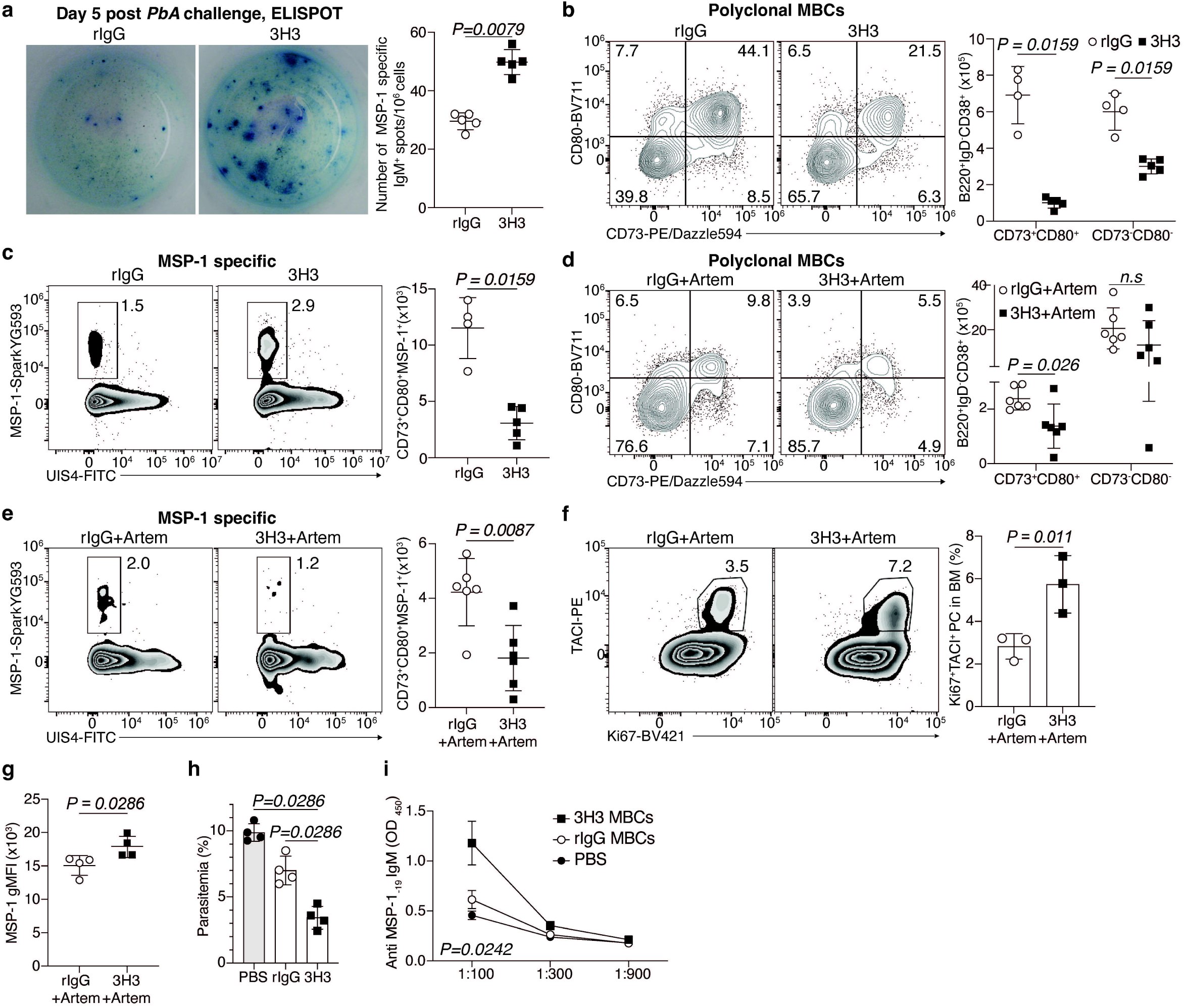
Enhanced recall potential and not abundance of MBCs drives protection following 4-1BB stimulation. **a,** Anti-MSP-1_-19_ IgM ELISPOT of bone marrow plasma cells (BMPCs) on 5 dpi post *PbA* challenge. Data are representative of *n=2* biologically independent experiments using *n=4* mice/group. **b-e**, *Py*-infected C57BL/6 mice were treated with 3H3 or isotype control as previously described. **d-f,** Mice were additionally treated with the antimalarial drug artemether at regular intervals as detailed elsewhere. Representative flow plots (left) and total numbers (right) of polyclonal (**b,d**) and MSP-1 specific (**c,e**) MBCs in the spleen. Data are either representative (**b,c**) or are pooled from *n=2* biologically independent experiments using *n=3-4* mice/group. **f,** Representative flow plots (left) and frequency (right) of proliferating BMPCs as detected by Ki67 and TACI staining. **g,** Geometric mean fluorescence intensity (gMFI) of MSP-1 staining in MSP-1 specific MBCs. Data are representative of *n=2* biologically independent experiments using *n=3* mice/group. **h,** Total MBCs from rIgG-and 3H3-treated mice were flow-sorted and transferred to μMT recipients that were infected with *Py* the following day. **h**, Summary graphs showing parasite burden on 12 dpi and **i**, anti-MSP-1_-19_ IgM serum antibody titers on 4 dpi. Data are representative of *n=2* biologically independent experiments using *n=4* mice/group. Data were analyzed by two-tailed Mann-Whitney U tests (**a-e** and **g-I)**, and Student’s *t* test (**f**).

Given that chemoprophylaxis using artemether preserved the protective effect of 3H3 (**Fig. 2e and f**), we asked whether the PC and MBC compartments showed similar trends to what was observed without artemether. To this end, mice were infected and treated as shown in **Fig. 2d** and splenic MBCs before *PbA* challenge as well as PCs in the bone marrow 5 days post challenge were characterized. We observed lower number of polyclonal (**Fig. 3d**) as well as the MSP-1 specific (**Fig. 3e**) CD73^+^CD80^+^ MBCs in 3H3-treated mice. Despite the reduced number of MBCs and in agreement with the survival data following *PbA* challenge (**Fig. 2e**), we observed a larger frequency of PCs that stained positive for Ki67 and TACI, indicative of a high proliferative burst (**Fig. 3f**). Taken together, these data suggest that, MBCs from 3H3-treated mice are qualitatively different and are capable to differentiate and proliferate into larger number of PCs upon antigen encounter. Since artemether treatment preserves the survival advantage and cellular phenotypes, we decided to adopt this regimen as a standard approach in future experiments. This will not only allow the mice to reach convalescence early (35-40 dpi with artemether compared to 85-90 dpi without), but will also simulate the chemoprophylactic approach often used in malaria-endemic regions.

Confirming our prediction that MBCs from 3H3-treated mice may be functionally superior, they showed a higher binding affinity to MSP-1 as evidenced by a higher geometric mean fluorescence intensity of MSP-1 staining (**Fig. 3g**). To formally compare the functional capacity of these MBCs upon antigen re-exposure, we flow-sorted MBCs from 3H3-and rIgG-treated mice and adoptively transferred them into separate groups of B cell deficient (μMT) recipients that were then infected with *Py* the following day. *Py*-infected μMT mice that received MBCs from 3H3-treated donors exhibited lower parasite load on 12 dpi compared to those receiving MBCs from rIgG-treated donors (**Fig. 3h**) and this correlated with higher serum antibody titers (**Fig. 3i**) on 4 dpi. Collectively, our data so far show that MBCs derived from 3H3-treated mice exhibit intrinsically superior recall potential and capacity to differentiate into antibody secreting PCs.

### Exogenous 4-1BB ligation drives extrafollicular MBCs

To understand the genetic basis for the functional differences between MBCs from 3H3-and rIgG-treated mice, we performed scRNA-, CITE-and V(D)J-sequencing on flow-sorted polyclonal (B220^+^IgD^-^GL7^-^CD38^+^) and MSP-1 specific (B220^+^IgD^-^GL7^-^CD38^+^Decoy^-^MSP1^+^) MBCs (**Supp Fig. 4a**) on day 38 post *Py* infection from both groups as detailed in **Fig. 4a**. As internal controls to verify our workflow, resting B cells (B220^+^IgD^+^) from either group were also spiked in to represent ∼ 25% of all cells sequenced in their respective polyclonal fraction. As shown in the UMAP (**Supp Fig. 4b**), the polyclonal MBCs grouped into 10 different clusters with the spiked-in resting B cells (IgD^+^) grouping separately in cluster 1 and representing roughly 25-27% of all cells sequenced (**Supp Fig. 4c**). To better highlight the potential differences among different MBC populations, polyclonal cells were re-clustered without the IgD^+^ population (**Fig. 4b**) into 9 clusters. Notably, cluster 0 was significantly more represented in the 3H3 group, comprising 55% of all MBCs compared to 28% in the rIgG group (**Fig. 4c**). This cluster exhibited increased expression of molecules involved with B cell activation (*Cd79a*, *Cd79b* and *Cd40*)^51–53^ and survival (*Tnfrsf13b* and *Tnfrsf13c*)^54–57^ (**Fig. 4d**). Cluster 2 was equally represented in both groups (19-20%) (**Fig 4c**) and expressed genes associated with robust activation and recall potential such as *Fcrl5*^58^ and *Il9r*^59^, respectively (**Fig. 4d**). In contrast, clusters 1 and 3 were reduced in the 3H3 group relative to the rIgG group (**Fig. 4c**) and were characterized by markers associated with stemness (*Sox4*, *Myb*, *Il7r* and *Cd93*)^60–63^ and genes associated with GC-derived MBCs (*Plxnb2*, *Basp1*, *Scimp* and *Tox*)^64^, respectively (**Fig. 4d**). Intriguingly, a closer look revealed that clusters 0 and 2, comprising around 75% of cells derived from 3H3-treated mice (**Fig. 4c**), exhibited a gene signature (**Supp table. 1**) indicative of an extrafollicular (EF) origin^64^ (**Fig. 4e and f**). This is consistent with our data showing a higher recall potential of MBCs from 3H3-treated mice, as EF-derived MBCs have been shown to be more proliferative and result in larger number of PCs^65^. Finally, clusters 4-8 that showed lower EF-derived signature comprised only 6% of MBCs in the 3H3 group compared to nearly 22% (**Fig. 4c**) in the rIgG group and contained mostly atypical B cells (**Fig. 4d, e and f**).

**Fig. 4:**
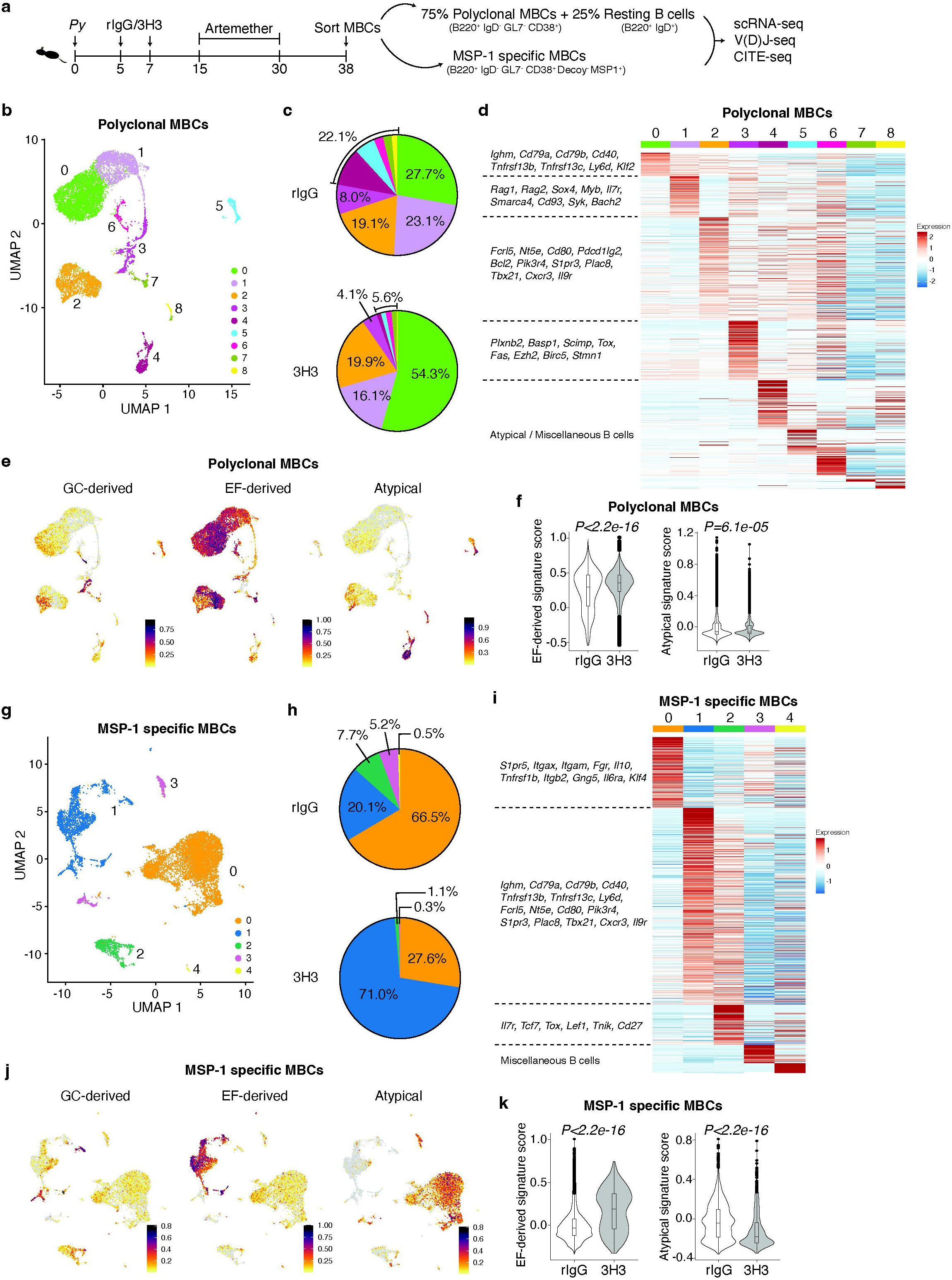
Exogenous 4-1BB ligation induces MBCs with an extrafollicular gene signature. **a,** Experimental design. C57BL/6 mice were infected and treated as indicated. Polyclonal and MSP-1 specific MBCs were flow-sorted and scRNA-seq, V(D)J sequencing and CITE-seq were performed. **b,** UMAP showing the identified clusters of polyclonal MBCs. **c,** Pie diagram depicting the frequency of respective clusters in the polyclonal MBC pool derived from rIgG and 3H3-treated mice. **d,** Heatmap depicting markers from each cluster with selected genes highlighted on the left. **e,f,** Cells were scored based on gene signatures of MBCs derived from a GC response, EF response or from atypical B cells and were visualized per cluster (**e**) and the prevalence of each of these signatures are compared between treatments as shown (**f**). **g,** UMAP showing clusters in MSP-1 specific MBCs. **h,** Frequency of clusters in MSP-1 specific MBCs from rIgG or 3H3-treated mice cells are shown in the pie chart. **i,** Heatmap of markers from each cluster with selected genes highlighted on the left. **j,k,** Cells were scored based on gene signatures of MBCs derived from a GC response, EF response or atypical B cells and visualized per cluster (**j**) and between treatments (**k**). Data from **f** and **k** were analyzed by Wilcoxon test. Cells were pooled from 3 mice per group prior to sequencing.

To further investigate if 3H3 treatment programs antigen-specific MBCs in a similar fashion, we analyzed sequences from MSP-1 specific cells. As shown in the UMAP (**Fig. 4g**), we identified 5 clusters, with cluster 0 being overrepresented in the rIgG group (67%), compared to only 28% in the 3H3 group (**Fig. 4h**) and marked by high expression of genes characteristic of atypical MBCs, such as *Itgax*, *Itgam*, *Itgb2* and *Fgr*^66,67^ (**Fig. 4i**). Conversely, cluster 1 comprised 71% of MSP-1 specific MBCs in the 3H3 group, but only 20% in the rIgG group (**Fig. 4h**) and characterized by a combination of markers present in clusters 0 and 2 of the polyclonal pool (**Fig. 4d and Fig. 4i**). Cluster 2 was less represented in the 3H3 group (1%) compared to the rIgG group (8%) and expressed markers associated with stemness, such as *Il7r*^68,69^ and *Lef1*^70^ (**Fig 4h and Fig 4i**). Clusters 3 and 4 made up less than 6% in the rIgG group and less than 0.5% in the 3H3 group (**Fig 4h**). Much like clusters 0 and 2 in the polyclonal pool (**Fig. 4e**), cluster 1 of the MSP-1 specific pool was enriched for an EF-derived signature (**Fig 4j**), which was more pronounced in 3H3-group (**Fig. 4k**), while cluster 0 (MSP-1 specific) from the rIgG group showed the highest atypical signature (**Fig. 4j and k**). These data collectively demonstrate that MBCs generated in mice following systemic 4-1BB stimulation is predominantly of EF origin, whereas those from control mice resemble atypical MBCs.

### 3H3 treatment generates MBCs with a distinct transcriptional profile supporting enhanced recall

Characterization of the major clusters revealed differences in the functional composition of the polyclonal and MSP-1 specific MBC pools from rIgG-and 3H3-treated mice. Importantly, cluster 0 in the polyclonal pool expressed the highest level of IgM (*Ighm*) (**Fig. 5a and Supp Fig. 5a**), a hallmark of a highly plastic and rapidly responding MBCs following *Plasmodium* rechallenge^65^; while cluster 2 co-expressed CD73 (*Nt5e*), CD80 (*Cd80*) (**Fig. 5b and Supp Fig. 5b**) and *Pdcd1lg2* (**Supp Fig. 5c**), markers indicative of superior recall^50^. Utilizing pseudotime inference using cluster 1 as the origin, the IgM^+^ population (cluster 0) appeared to be in the early stage of its differentiation process, while clusters 2, 4 and 5 were closer to being terminally differentiated (**Fig. 5c**). In contrast to MBCs from rIgG-treated mice (blue arrows, **Fig. 5c**), polyclonal MBCs from 3H3-treated mice favored the early and plastic stage^65^ (cluster 0, IgM^+^) (thick red arrow, **Fig. 5c**) and had their differentiation skewed away from the terminal atypical trajectories (thin red arrows, **Fig. 5c**). This will result in the preferential accumulation of IgM^+^ EF MBCs and reduction of atypical MBCs in the 3H3 group (**Fig. 4c**). Characterization of MSP-1 specific pool showed that cluster 1 from 3H3-treated mice co-expressed high levels of IgM (*Ighm*) (**Fig. 5d and Supp Fig. 5d**), CD73 (*Nt5e*), CD80 (*Cd80*) (**Fig. 5e and Supp Fig. 5e**) and *Pdcd1lg2* (**Supp Fig. 5f**). Pseudotime analysis using cluster 2 as the origin showed that MBCs from 3H3-treated mice shifted their differentiation towards the EF trajectory (thick red arrow, **Fig. 5f**) and away from the atypical trajectory (thin red arrow, **Fig. 5f**). While MBCs from the rIgG group showed a higher propensity to take the atypical trajectory, away from the EF route (blue arrow, **Fig. 5f**). The data so far showing that MBCs from 3H3-treated mice are more likely to have an EF origin is further corroborated by our V(D)J sequencing data showing they have significantly lower mutational frequency in their BCRs (**Supp Fig. 5g and h**).

**Fig. 5:**
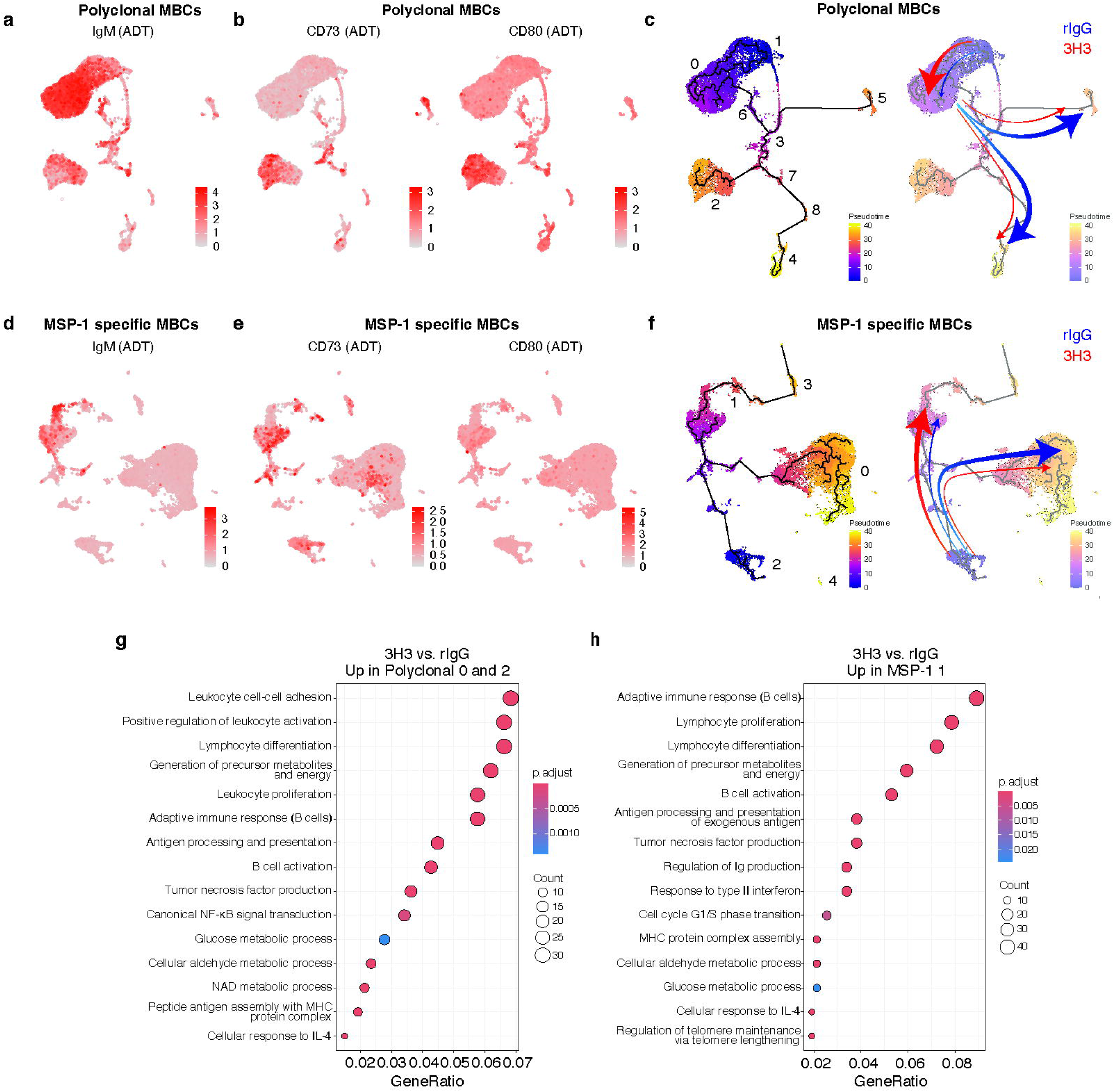
4-1BB stimulation skews MBC differentiation towards a highly functional population. C57BL/6 mice were infected and treated as indicated in Fig. 4a. Distribution of IgM (**a**), CD73 and CD80 (**b**) expression among the polyclonal MBCs clusters. **c,** Pseudo time trajectories (left) and their representative contributions towards each cluster in polyclonal MBCs in rIgG-and 3H3-treated mice (right) based on their frequencies in Fig. 4c. Distribution of IgM (**d**), CD73 and CD80 (**e**) expression among the MSP-1 specific MBCs clusters. **f,** Pseudo time trajectories (left) and their representative contributions in MSP-1 specific MBCs in rIgG-and 3H3-treated mice (right) based on their frequencies in Fig. 4h. Gene ontology (GO) analysis of upregulated genes in MBCs from 3H3-treated mice compared to MBCs from rIgG-treated mice in clusters 0 and 2 of polyclonal MBCs combined (**g**) and cluster 1 of MSP-1 specific MBCs (**h**).

To explore functional differences within the EF clusters, we analyzed differentially expressed genes (DEGs) between rIgG and 3H3 in clusters 0 (IgM^+^) and 2 (CD73^+^CD80^+^) from the polyclonal pool as well as cluster 1 from the MSP-1 specific pool **(Supp. Table 2)**. Notably, 122 DEGs were upregulated across all 3 clusters (**Supp**. Fig 5i). Approximately 88% of polyclonal cluster 2 DEGs were shared with polyclonal cluster 0 and/or MSP1^+^ cluster 1, with only 21 unique genes. In addition, more than half of the DEGs in polyclonal cluster 0 and MSP-1 specific cluster 1 were shared, underscoring the similarities between these two clusters (**Supp**. Fig 5i). Given that most DEGs in polyclonal cluster 2 overlapped with the other 2 clusters (**Supp**. Fig 5i), we further investigated the biological processes regulated by the combined DEGs in polyclonal clusters 0 and 2 (**Fig. 5g**) and MSP-1 specific cluster 1 (**Fig. 5h**). Notably, both polyclonal and MSP-1 specific MBCs from 3H3 group showed increased expression of genes associated with heightened capacity for activation, proliferation and antigen presentation as well as certain metabolic pathways, possibly to meet the energetic and biomass demands to support these processes (**Fig 5g and Fig 5h**). In addition, MBCs from 3H3-treated mice also showed enhanced cellular response to IL-4, a cytokine shown to downregulate GC related genes and enhance MBC generation and activation^71,72^, which may enforce the MBCs to take the extrafollicular route. Aligning with these data, both the polyclonal (clusters 0 and 2, **Supp Fig. 5j**) and MSP-1 specific MBCs (cluster 1, **Supp Fig. 5k**) also downregulated genes involved in pathways related to increased apoptosis, reduced transcription factor binding, gene silencing and metabolism among others. Collectively, these data confirm that exogenous ligation of 4-1BB results in the generation of MBCs with an EF origin that are, although numerically fewer, functionally superior.

### 3H3-treatment induced protection is dependent on IFNγ and B cell intrinsic T-BET:IL-9R axis

Previous studies have highlighted the role of B cell intrinsic IFNγ signaling and T-BET expression ^73–75^ for enhanced MBC recall and PC differentiation. Importantly, exogenous stimulation of 4-1BB was shown to result in higher levels of IFNγ secretion by CD4 T cells^76^. When splenocytes were stimulated in the presence of PMA/Ionomycin and brefeldin A, we observed that CD4 T cells derived from 3H3-treated mice were indeed capable of secreting higher levels of IFNγ (**Supp Fig 6a**). Additionally, B cells cultured *ex vivo* with IFNγ showed a significant increase in T-BET expression within 18 hours (**Supp Fig. 6b**). In line with this, we also observed significantly higher expression of T-BET in activated (IgD^-^) B cells derived from 3H3-treated mice around 15 dpi (**Supp Fig. 6c**). T-BET expression in B cells during the effector phase was shown to enhance protection from rechallenge by mediating class-switching to IgG2c antibody isotype^73^. Consistent with this finding, we observed higher increases in IgG2c levels following *PbA* challenge in 3H3-treated mice (**Supp Fig. 6d**). Given these data, we asked whether 3H3-mediated superior protection is IFNγ dependent. To test this, mice were infected with *Py* and treated with 3H3 or rIgG and IFNγ was blocked using IFNγ blocking antibody (XMG1.2) on indicated timepoints. Mice were then challenged with *PbA* (around 90 dpi) and survival was monitored (**Supp Fig. 6e**). As expected, 3H3-treated mice lost protection following *PbA* challenge when IFNγ was blocked (**Supp Fig. 6f**). Given the link between IFNγ and T-BET, we then asked whether B cell intrinsic T-BET expression is necessary for 3H3-mediated protection. To this end, we generated mixed bone marrow chimeras in which T-BET deficiency was restricted to the B cell compartment (B*^Tbx^*^21-/-^) (**Supp Fig. 6g**) and infected them with *Py,* followed by 3H3 treatment. Similar to our results from XMG1.2 treatment, *PbA* challenge of convalescent B*^Tbx21^*^-/-^ mice showed significantly reduced survival compared to B^WT^ mice (**Supp Fig. 6h**). Taken together, these highlight the importance of IFNγ:T-BET axis during *Plasmodium* infection and show that the enhanced protection observed in mice subjected to exogenous 4-1BB ligation is dependent on this axis.

IL-9 receptor (IL-9R) signaling on MBCs has also been shown to result in superior recall^59^. In line with this, MSP-1 specific MBCs from 3H3-treated mice subjected to IFNγ blockade and B*^Tbx21^*^-/-^ mice showed significantly lower levels of IL-9R expression (**Supp Fig. 6i and j**) suggesting that B cell intrinsic IFNγ:T-BET axis is key to this phenomenon. To investigate the contribution of IL-9R to higher MBC recall, we searched for distinct *Il9r*-expressing cluster(s) in our scRNA-seq data and identified polyclonal (cluster 2) (**Fig. 6a**) and MSP-1 specific (cluster 1) (**Fig. 6b**) which also co-expressed other signature MBC markers such *as Nt5e, Cd80, Pdcd1lg2* among others (**Fig. 4d**). Quantitative real-time PCR analysis on flow-sorted MBCs from 3H3-treated mice also showed significantly higher levels of *Il9r* expression compared to those from rIgG-treated mice (**Supp.** Fig 7a). In an attempt to draw parallels among populations from our transcriptomic and flow cytometry datasets, we clustered our multi-parameter flow cytometry data in a UMAP space (polyclonal, **Fig. 6c** and MSP-1 specific, **Fig. 6f**). As shown in **Fig. 6c and d**, the polyclonal pool clustered into 9 populations (*A-I*), of which populations *D* and *I* denoted the MSP-1 specific pool. While populations *E, F* and *I* expressed IL-9R, population E failed to express key functional markers such as CD73 and CD80 making it less likely to be involved in an immediate recall following antigen re-exposure (**Fig. 6d** and **Supp Fig. 7b**). Importantly and in agreement with this, we observed that the CD73^+^CD80^+^ MBC population from 3H3-treated mice expressed significantly higher levels of IL-9R compared to controls (**Fig. 6e**). These data also suggested that population *I* may be functionally similar to population *F* and together they may represent antigen-specific MBC pools that are swiftly recalled following antigen-reencounter. Phenotypic dissimilarity of populations *D* and *I* also highlights the functional heterogeneity within a given antigen-specific pool. This was confirmed by further clustering the MSP-1 specific pool which yielded 5 different populations, mostly differing in their expression of key functional MBC markers (**Fig. 6f and g**). As with the polyclonal cells (**Fig.6e**), MSP-1 specific MBCs from 3H3-treated mice also expressed higher levels of IL-9R (**Fig. 6h**) suggestive of their superior recall potential. Given the parallels between the gene and surface maker expression data, polyclonal cluster 2 may be identical to population *F* and they may represent MBCs equipped to readily engage in a recall response. Given the dependency on IFNγ and B cell intrinsic T-BET expression for heightened survival of 3H3-treated mice, we hypothesized that IL-9R expression may also be regulated by this axis. To test this, B cells from naïve mice were activated *ex vivo* in the presence or absence of IFNγ and IL-9R expression was evaluated after 3 days in culture. Although activation of B cells alone resulted in increase in IL-9R expression, it was further increased when IFNγ was added to the culture (**Supp Fig. 7c**).

**Fig. 6:**
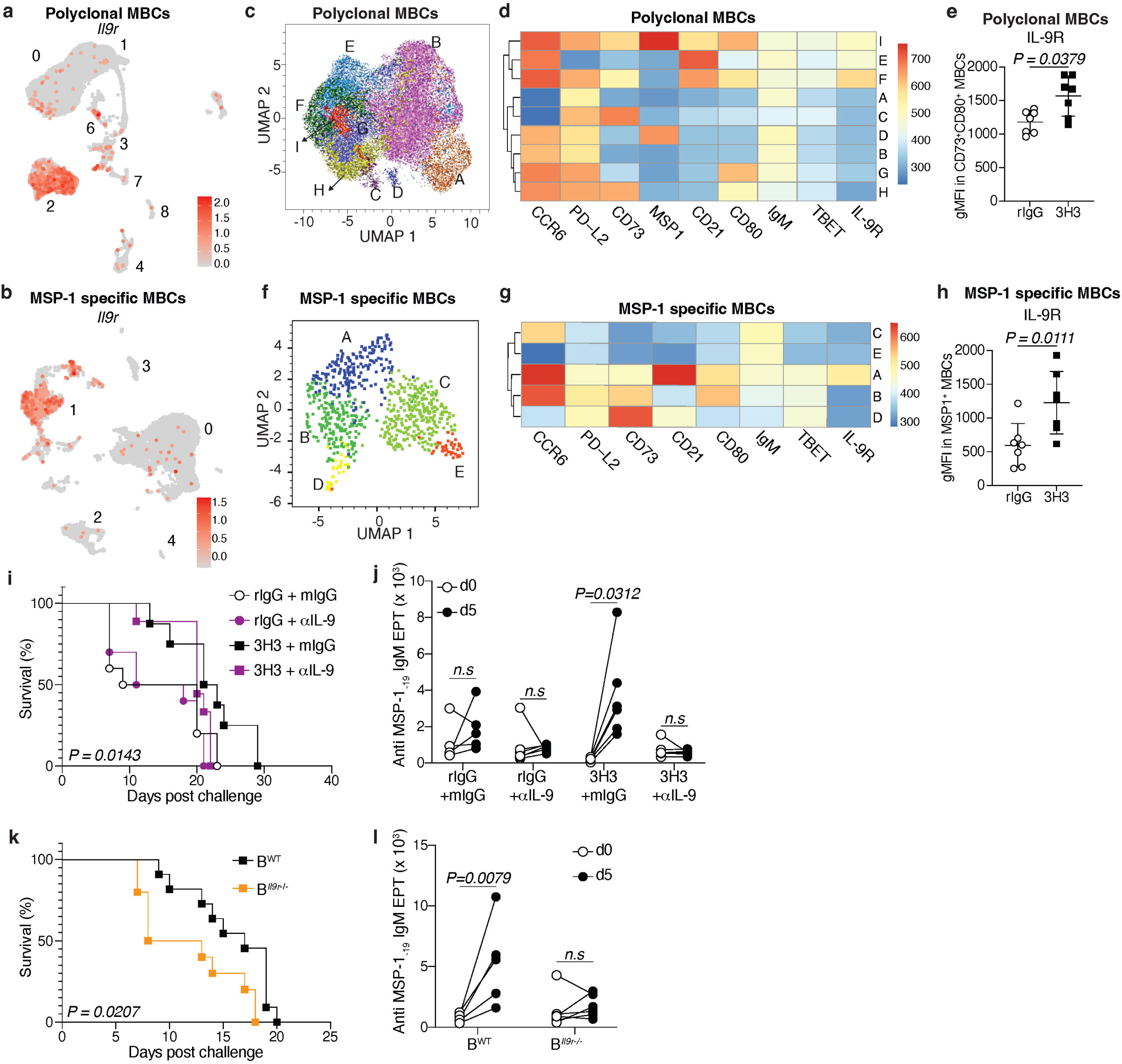
3H3-driven protection is dependent on IL-9:IL-9R signaling. C57BL/6 mice were infected and treated as indicated in Fig. 4a. UMAP clustering of scRNA-seq data showing *Il9r* expression among polyclonal (**a**) and MSP-1 specific MBC clusters (**b**). **c,** UMAP clustering of polyclonal MBCs on 38 dpi generated by FlowSOM. **d,** Expression levels of various proteins using FlowSOM clustering of polyclonal MBCs. **e,** Geometric mean fluorescence intensity (gMFI) of IL-9R in polyclonal MBCs on 38 dpi. Data is pooled from *n=2* biologically independent experiments using n=4 mice/group/experiment. **f,** UMAP clustering of MSP-1 specific MBCs on 38 dpi generated by FlowSOM. **g,** Expression levels of various proteins using FlowSOM clustering of MSP-1 specific MBCs. **h,** gMFI of IL-9R in MSP-1 specific MBCs on 38 dpi. Data is pooled from *n=2* biologically independent experiments using n=4 mice/group/experiment. **i,j,** WT mice were infected with *Py* and treated with rIgG or 3H3 and then with the antimalarial drug artemether at regular intervals. One day prior and after *PbA* rechallenge, mice were treated with αIL-9 neutralizing antibody or isotype control (mIgG) (as detailed in **Supp Fig. 7d**). Survival was monitored (**i**) and fold change in anti-MSP-1_-19_ IgM serum antibody EPT between 0 and 5 dpi was assessed (**j**). Data is pooled from *n=2* biologically independent experiments using *n=6* mice/group. **k**,**l,** B cell-specific WT (B^WT^) and IL-9R KO (B^IL-9R KO^) chimeras generated and were infected with *Py* and treated with 3H3 on days 5 and 7 post infection and then with the antimalarial drug artemether at regular intervals (**Supp Fig.7e**). On 38 dpi, during the convalescent phase, mice were rechallenged with *PbA*. Survival was monitored (**k**) and fold change in anti-MSP-1_-19_ IgM serum antibody EPT between 0 and 5 dpi was assessed (**l**). Data is pooled from *n=2* biologically independent experiments using *n=11* mice/group (**k**) and are representative of *n=2* biologically independent experiments using *n=5-6* mice/group (**l**). For **e**, **h**, **j** and **l** two-tailed Mann-Whitney U tests were used for statistical analysis. For **i** and **k**, data were analyzed by Mantel-Cox.

To investigate whether the enhanced protection following *PbA* challenge in 3H3-treated mice is dependent on IL-9:IL-9R signaling, we blocked IL-9 in convalescent mice prior to *PbA* challenge and monitored survival (**Supp Fig. 7d)**. As shown in **Fig**. **6i**, the enhanced protection observed in 3H3-treated mice was reversed in mice treated with IL-9 blocking antibody compared to controls; rIgG-treated mice did not show any difference in survival. Supporting our observation, 3H3-treated mice subjected to IL-9 neutralization exhibited a blunted antibody recall following *PbA* challenge (**Fig. 6j**), indicating that IL-9 signaling is important for 3H3-mediated protection. Next, we interrogated whether B cell-intrinsic IL-9R expression is important for the enhanced protection following 3H3 treatment. To accomplish this, we generated competitive mixed bone marrow chimeras in which IL-9R deficiency is restricted to B cells (B*^Il9r-/-^*) (**Supp Fig. 7e**). Since we did not see any difference in the rIgG treated mice following IL-9 blockade (**Fig. 6i**), we confined our experiment to 3H3-treatment in B*^Il9r-/-^* and B^WT^ mice. On 38 dpi, mice were challenged with *PbA* and survival was monitored. Notably, we observed that B*^Il9r-/-^* mice showed significantly shorter survival than B^WT^ mice following *PbA* challenge (**Fig. 6k**) and a dampened MBC recall response as evident from the lower fold change in serum antibody titers (**Fig. 6l**). These data show that the MBC recall following *PbA* challenge in 3H3-treated mice is dependent on the B cell-intrinsic IL-9R expression. Collectively, our data highlight a previously underappreciated IFNγ:T-BET:IL-9R axis present in MBCs following *Plasmodium* infection, which could potentially be modulated to enhance humoral immune memory.

### Exogenous 4-1BB ligation poises MBCs for swift recall by altering their epigenome

Given that both polyclonal and MSP-1 specific MBCs from 3H3-treated mice downregulated genes associated with transcriptional silencing and regulation of transcription factor activity (**Supp.** Fig 5j and k), we wanted to determine whether these MBCs are imprinted with an epigenetic signature supporting superior recall. To this end, we flow-sorted polyclonal and MSP-1 specific MBCs from 3H3-or rIgG-treated mice (**Fig. 7a**) as before and performed scATAC-seq. Initial clustering based on transposase activity populated 4 different clusters in the MSP-1 specific cells (**Supp.** Fig 8a) and 6 clusters in the polyclonal pool (**Supp.** Fig 8b). Given the importance of B cell-intrinsic IL-9R expression in mice treated with 3H3, we wanted to identify the IL-9R expressing population in our scATAC-seq dataset and examine the accessibility of genes that can potentiate PC differentiation. Thus, we integrated our scRNA-seq (from **Fig. 4 and 5**) with the scATAC-seq datasets and used the transferred labels to annotate the ATAC clusters from MSP1-specific (**Fig. 7b**) and polyclonal MBCs (**Supp Fig. 8c**). As shown in **Fig. 7b**, following integration, most cells fell in either cluster 0 or 1. Since cluster 1 in MSP-1 specific MBCs expressed the highest level of *Il9r* (**Fig. 6b**), we examined whether the *Il9r* gene locus in this cluster is still accessible. As shown in **Fig. 7c**, we observed significantly higher accessibility in the *Il9r* locus in cluster 1 compared to cluster 0, showing that MBCs in cluster 1 are imprinted with an epigenome that favors higher *Il9r* expression. In line with our prediction that IL-9R expressing MBCs are primed to quickly differentiate into PCs, cluster 1 also showed enhanced accessibility for *Prdm1* locus (**Fig. 7d**) as well as for key transcription factors (TF) binding motifs (XBP1, ATF3, MYC, ATF6, KLF6 and BHLHE40) (**Fig. 7e**) associated with enhanced PC differentiation. Importantly and similar to cluster 1 in the MSP-1 specific pool, cluster 2 in the polyclonal pool also showed significantly enhanced *Il9r* locus accessibility (**Supp Fig. 8d**) as well as markers of higher activation potential (*Fclr5*, **Supp Fig. 8e**), enhanced PC differentiation (*Atf6*, **Supp.** Fig 8f) and antiapoptotic signaling (*Bcl2*, **Supp Fig. 8g**). In addition, TF binding motifs important in PC differentiation (KLF4, ATF3, XBP1 and IRF4) as well as cell proliferation (MYC) and IFNγ signaling (IRF1) were also accessible in cluster 2 from the polyclonal pool (**Supp Fig.8h**). Given that *Il9r* expression correlates with enhanced protection in 3H3-treated mice, we compared the chromatin accessibility of *Il9r*-expressing MBCs between 3H3-and rIgG-treated groups. In strong agreement with our survival and phenotypic data, MBCs from 3H3-treated mice in cluster 1 not only showed continued accessibility for *Il9r* locus (**Fig. 7f**), but were also enriched for the key transcription factor binding motifs associated with enhanced PC differentiation and proliferation (**Fig. 7g**). Notably, we also observed that T-BET and IRF1 binding motifs also showed higher accessibility further reinforcing our finding that the superior recall we observe in 3H3-treated mice is dependent on IFNγ:T-BET:IL-9R axis.

**Fig. 7:**
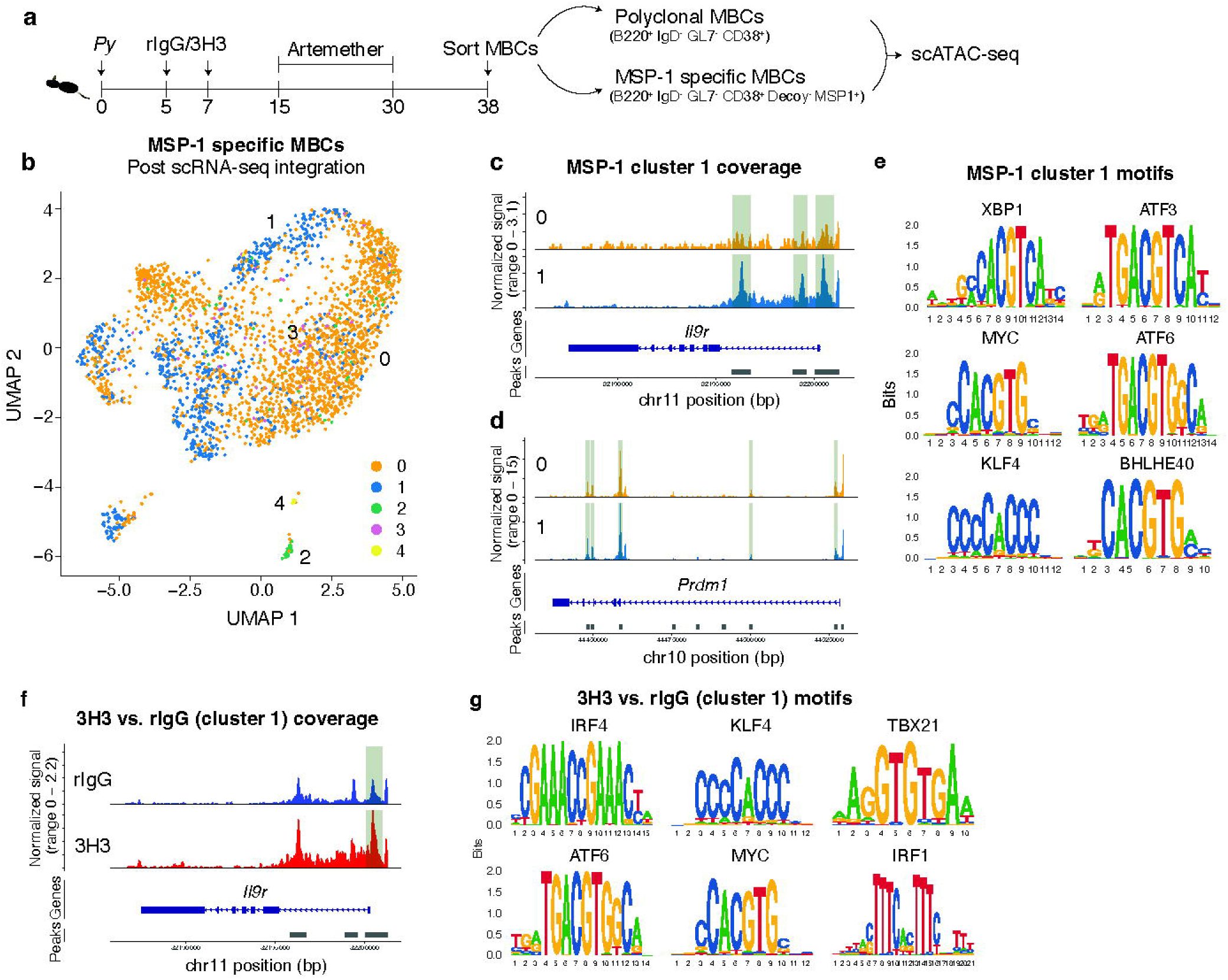
Altered chromatin landscape in MBCs following 4-1BB stimulation poises them to readily become PCs upon recall. **a,** Experimental design. C57BL/6 mice were infected and treated as indicated. Polyclonal and MSP-1 specific MBCs were flow-sorted and scATAC-seq was performed. **b,** UMAP clustering of MSP-1 specific MBCs using scATAC-seq data with transferred labels from scRNA-seq post data integration. Comparison of *Il9r* (**c**) and *Prdm1* (**d**) accessibility between clusters 0 and 1 of MSP-1 specific MBCs. **e,** Motif analysis of differentially accessible regions (DARs) in cluster 1 of MSP-1 specific MBCs compared to the other clusters. **f,** Comparison of *Il9r* accessibility in cluster 1 between MSP-1 specific MBCs obtained from rIgG and 3H3-treated mice. **g,** Motif analysis of differentially accessible regions (DARs) in 3H3 compared to rIgG in cluster 1 of MSP-1 specific MBCs. Sequenced nuclei were pooled from 4 mice per group.

## Discussion

In this study, we investigated the effects of exogenous 4-1BB ligation and observed an unexpected outcome in MBC generation in an experimental mouse model of malaria. Despite the disruption of the effector GC response and a total reduction in MBC numbers, the MBC recall response was significantly enhanced, providing protection upon reinfection. While our data draw some parallels with a previous study using a viral infection model ^77^ showing a derailed GC response, our findings on humoral immune memory response are unique. Single-cell transcriptomic analysis of the MBCs showed a predominant extrafollicular rather than a GC-derived signature, with IL-9R emerging as a key determinant for the enhanced recall. Using a combination of genetic, chimeric, and biochemical approaches, we identify a previously unappreciated axis comprising IFNγ and B cell intrinsic T-BET expression during the effector phase, potentially driving the expression of IL-9R. Sequencing for transposase-accessible chromatin regions in flow-sorted MBCs from *Plasmodium-*infected mice treated with exogenous 4-1BB agonist also revealed an epigenomic landscape that primed them for rapid differentiation into PCs.

Similar to our study, *Ehrlichia muris (E. muris)* infection alone without the need of any agonistic antibody to 4-1BB, resulted in a derailed GC response^78^. Notably, the MBCs that appeared in *E. muris* infection were CD11c^+^ and T-dependent but seldom expressed GC-related markers^79^. Thus, the antigen-specific IgM^+^ MBCs (cluster 1) in our study are largely similar to MBCs mentioned in the study above^79^, but express FCLR5 and not CD11c. Co-expression of CD11c, CD11b and FCLR5 are considered markers for atypical MBCs, with predominantly inhibitory functions ^80–82^. Based on the lack of CD11b and CD11c expression combined with the expression of various activation markers (**Fig. 4 and 6**) including FCLR5^58^, we contend that the IgM^+^ MBCs from 3H3-treated mice (cluster 1) are different from the inhibitory atypical MBCs.

IFNγ-driven T-BET expression was shown to be pivotal to increased protection following *Plasmodium* rechallenge^73^. Reliance on IFNγ:T-BET axis in 3H3-treated mice for enhanced recall, along with a pronounced EF signature of these MBCs, is in agreement with previous findings showing that B cell-intrinsic IFNγ signaling reinforces EF B cell responses^83^. Previous studies have also independently shown the role of IL-9R expression for enhanced recall^59^. 3H3-treated but not rIgG-treated mice, seem to depend on IL-9R expression specifically on B cells for protection, which may be explained by the far lower IL-9R expression in MBCs derived from rIgG-treated mice. Our scATAC-seq data shows that IRF1 binding motifs are readily accessible in *Il9r* expressing MBCs from both groups showing responsiveness to IFNγ^84^. More importantly, MBCs from 3H3-treated mice have a more pronounced accessibility to this region along with T-BET binding motifs. This further strengthens the possibility that IFNγ-driven T-BET may influence the expression of IL-9R in MBCs from 3H3-treated mice. We also observed lower IL-9R levels in B cells lacking T-BET expression or when IFNγ was blocked, which is complemented by our finding that IL-9R expression can be enhanced in B cells when subjected to IFNγ *ex vivo*. Together, these data suggest a potential role of IFNγ:T-BET axis in IL-9R expression and raises the possibility that T-BET levels may have to reach a specific threshold to enforce IL-9R expression. While these observations are B cell-centric, the source of IL-9 during the memory phase to signal via IL-9R remains to be investigated. As such, stimulation of various TNFRSF members have been shown to promote IL-9 secretion from CD4 T cells in multiple experimental settings^85,86–88^. Although our IL-9 neutralization experiments suggest the presence of an IL-9 source prevalent during the memory phase, a direct determination of the identity of the cellular source is yet to be done. IL-9 neutralization during the memory phase compromised survival and MBC recall only in 3H3-treated mice, signifying the role of IL-9:IL-9R signaling following 4-1BB stimulation. To our knowledge, our study remains the first of its kind to link the IFNγ:T-BET axis with IL-9R in MBC recall. Despite mining numerous publicly available human malaria datasets studying MBCs, we were unable to find a high IL-9R expressing population that could be correlated with enhanced protection upon pathogen re-exposure. This is due to the fact that none of these studies involved the use of immunomodulatory approaches that potentially drive high T-BET in B cells, which may be key to driving IL-9R. Indeed, our own *ex vivo* data with murine B cells stimulated with IFNγ resulted in the upregulation of T-BET first and IL-9R later (T-BET expression preceding IL-9R). A fate mapping approach capable of tracking T-BET expression would be the ideal strategy to directly ask this question, but it remains beyond the scope of this manuscript.

Although many studies have focused on the effect of 4-1BB on CD8 T cells^89,90^, systemic ligation of 4-1BB has been shown to reprogram CD4 T cells in a virus infection setting as well^76^. It is unlikely that 4-1BB stimulation on CD8 T cells may play a significant role in our studies, as CD8 T cell depletion in 3H3-treated mice still disrupted the effector germinal center (GC) response and resulted in substantial loss of parasite control, similar to observed in CD8 T cell replete WT mice (*Caloba and Vijay unpublished data*). B cells are also known to express 4-1BB^91^, but we did not explore the role of 4-1BB stimulation on B cells as CD4-specific 4-1BB^-/-^ bone marrow chimeras failed to respond to 3H3 treatment as it did not derail the parasite control. These findings indicate that in rodent malaria models, the effects of exogenous 4-1BB stimulation are dependent on CD4 T cell intrinsic 4-1BB expression.

Here we provide data showing that exogenous 4-1BB stimulation is capable of programing a predominantly EF B cell response that generates antigen-specific MBCs with superior recall potential. Importantly, although the MBCs in mice that received 4-1BB stimulation is at a numerical disadvantage, a large fraction of those cells possess a distinct epigenetic landscape and transcriptional signature poising them for swift and efficient recall. Single-cell trajectory analysis also reveals that MBCs from 3H3-treated mice have a tendency to stay more plastic within the extrafollicular space. Our data also brings up the argument that GC response during effector phase cannot always be a predictor for the ensuing memory response. We show that 4-1BB is a compelling target for host-directed immunotherapy in infections that are reliant on MBC-driven antibody response for protection. While the widescale use of a monoclonal antibody as an immunotherapy may be cost prohibitive, the prospect of synthesizing small molecule activators of 4-1BB for achieving potent humoral memory immune activation and thus durable and long-lasting protection may be practical.

## Supporting information

Supp Figures combined

## Acknowledgements

The authors thank all past and present members of the Vijay Lab for the helpful dicussions and technical assistance. We thank Drs. Fabio Re (RFUMS) and Danielle Stanisic (Griffith University) for the critical review of the manuscript. We thank the RFUMS flow cytometry facility for the instrumentation as well as the Biological Research Facility for animal husbandry and maintenance. We thank Dr. Stefan Green, Kevin Kunstman, Edith Perez and Ping Li at the Genomics and Microbiome Core Facility at Rush University for help with sequencing. This work was supported by the RFUMS startup funds and Schweppe Scholar Award to R.V and the AAI 2024 Careers in Immunology Fellowship to C.C. and R.V.

## Author Contributions

R.V. conceived the study. C.C. and R.V. designed, executed, analyzed and wrote the manuscript. C.C. also performed the bioinformatic data analyses. A.J.S. executed experiments. T.A.L., L.J., and A.R., provided technical assistance. A.M.M and S.E.L. generated and provided reagents, T.H.W. provided reagents as well as provided intellectual input, A.M.C. and M.H.K. generated and provided reagents, M.H.K. provided intellectual input, J.P.W. provided technical assistance for bioinformatic analyses.

