## Supplementary material for "Systemic 4-1BB stimulation augments extrafollicular memory B cell formation and recall responses during *Plasmodium* infection": Supp Figures combined

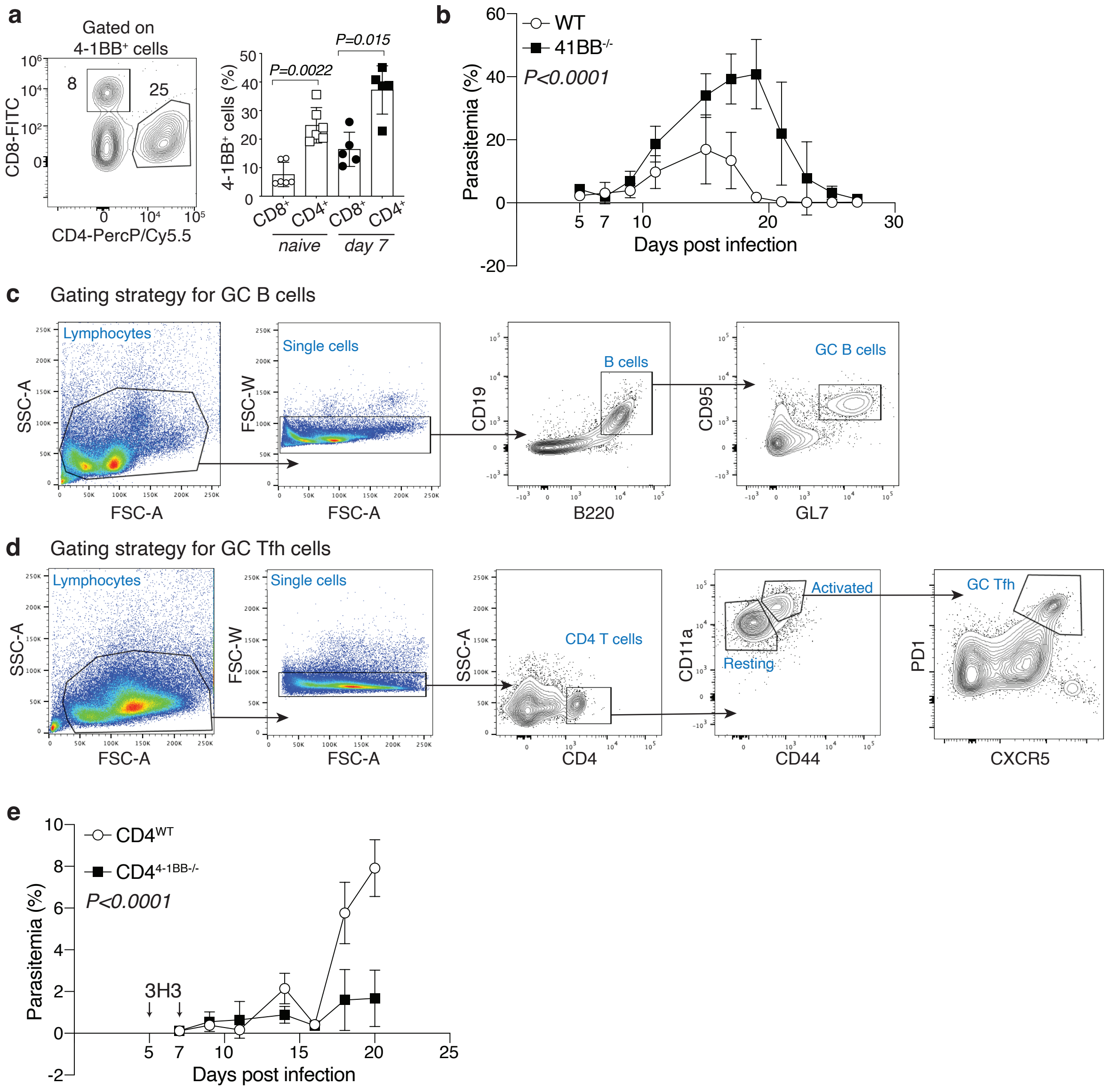


**Supp Fig. 1: 4-1BB is highly expressed in CD4 T cells during *Plasmodium* infection. a**, Frequency of 4-1BB expressing cells in CD8 and CD4 T cell pools. Data are pooled from 2 biologically independent experiments and analyzed by two-tailed Mann-Whitney U test. **b,** Kinetics of parasite burden in WT and 4-1BB$^{-/-}$ mice. Data are representative of 2 biologically independent experiments and analyzed by two-way ANOVA. **c,d,** Gating strategy for GC B cells (**c**) and Tfh cells (**d**). **e,** Kinetics of parasite burden in CD4 T cell specific 4-1BB$^{-/-}$ and WT competitive mixed bone marrow chimera treated with 3H3 on 5 and 7 dpi. Data are representative of 2 biologically independent experiments and analyzed by two-way ANOVA.

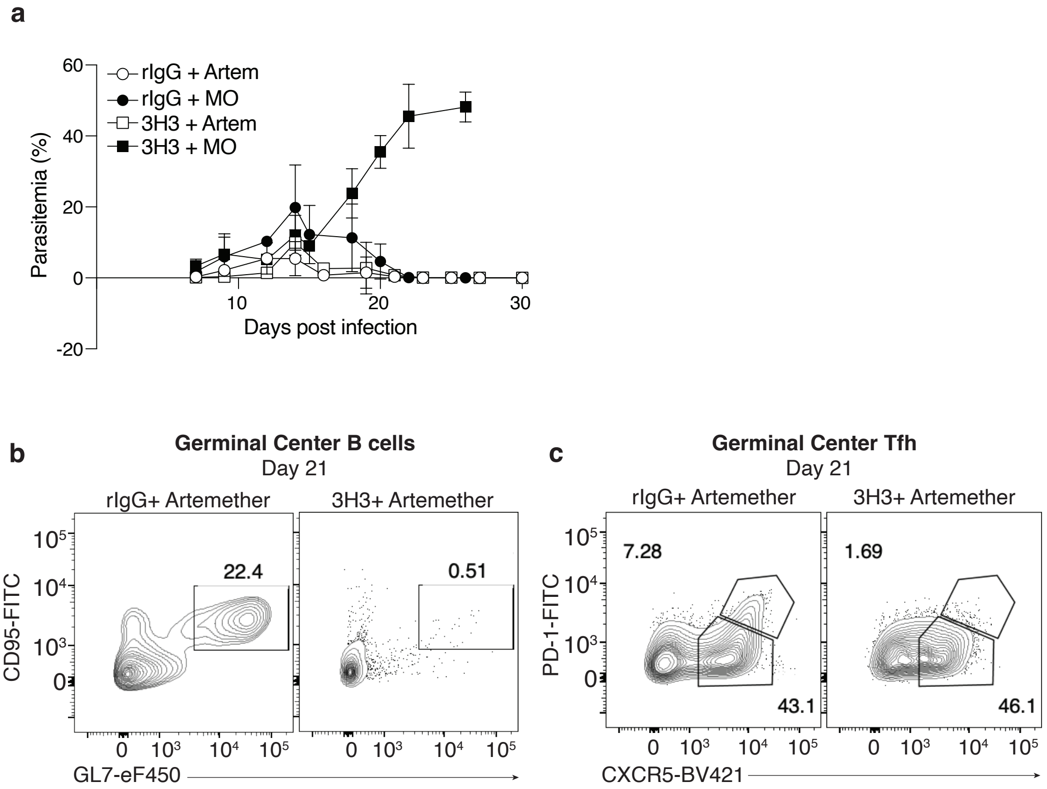


**Supp Fig. 2: 4-1BB stimulation derails GC response independently of antigen load. a**, Kinetics of parasite burden. Data are representative of at least 2 biologically independent experiments. **b,c,** Representative plots of GC B cells (**b**) and Tfh cells (**c**). Data are representative of at least 2 biologically independent experiments.

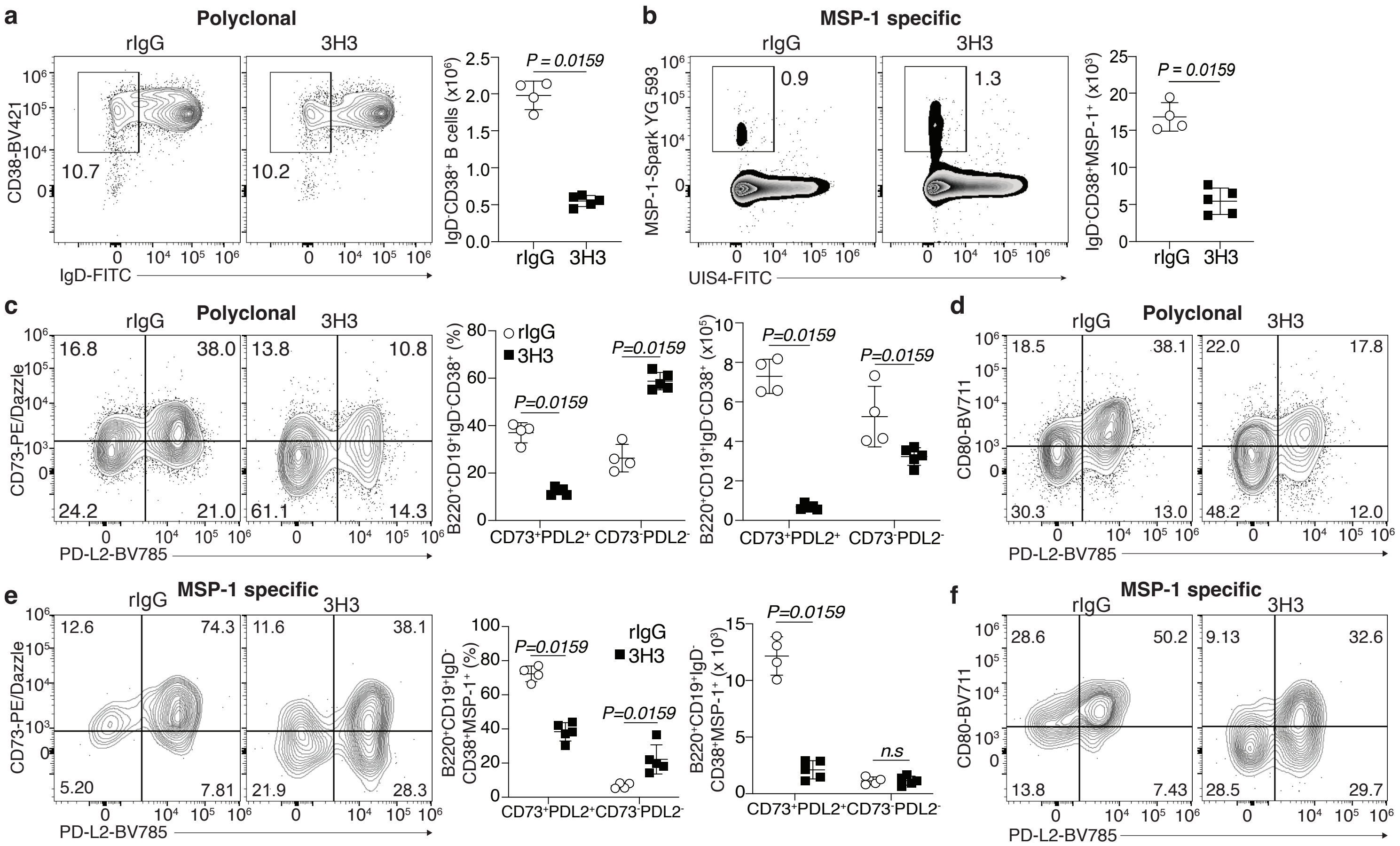


**Supp Fig. 3: Enhanced recall potential and not abundance of MBCs drives protection following 4-1BB stimulation. a,b,** Representative plots and total numbers of polyclonal (**a**) and MSP-1 specific (**b**) MBCs. Data are representative of 2 biologically independent experiments. **c,d,** Representative plots and total numbers of $CD73^{+}PD\text{-}L2^{+}$ (**c**) and $CD80^{+}$ $PD\text{-}L2^{+}$ (**d**) MBCs in the polyclonal pool. Data are representative of 2 biologically independent experiments. **e,f,** Representative plots and summary graphs of $CD73^{+}PD\text{-}L2^{+}$ (**e**) and plots of $CD80^{+}$ $PD\text{-}L2^{+}$ (**f**) MBCs in the MSP-1 specific pool. Data are representative of 2 biologically independent experiments. All data were analyzed by two-tailed Mann-Whitney U tests.

Supp Figure 4

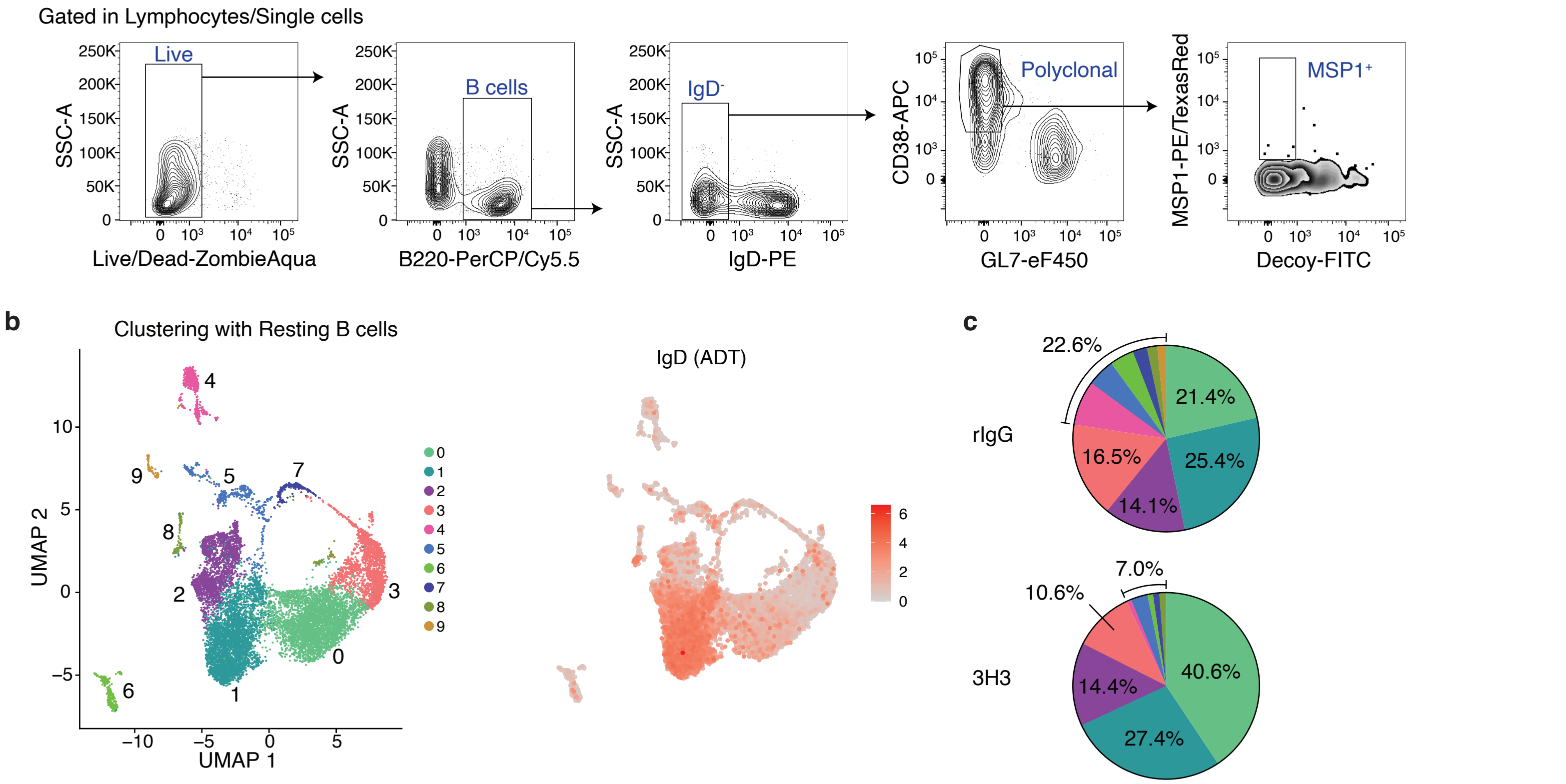


**Supp Fig. 4: Exogenous 4-1BB ligation induces MBCs with an extrafollicular gene signature. a,** Gating strategy for sorting of resting B cells (IgD⁺), polyclonal and MSP-1 specific (MSP-1⁺) MBCs. **b,** UMAP clustering (left) of MBCs containing the resting B cell population (right). **c,** Frequency of clusters in rIgG and 3H3 cells. Sequenced cells were pooled from 3 mice per group.

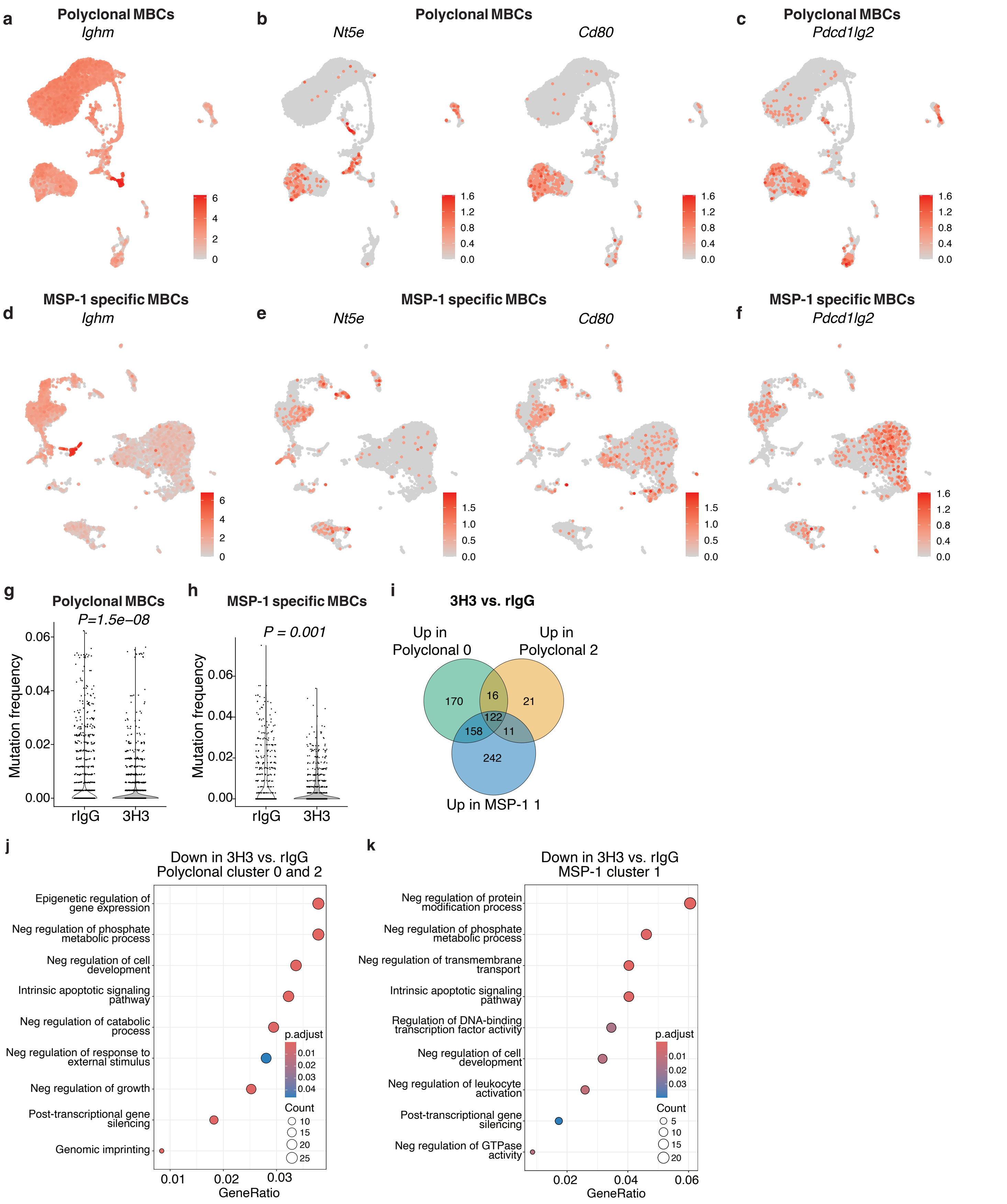


**Supp Fig. 5: 4-1BB stimulation skews MBC differentiation towards a highly functional population. a,b,c,** Distribution of *Ighm* (**a**), *Nt5e* (CD73) and *Cd80* (**b**) and *Pdcd1lg2* (PD-L2, **c**) expression among the polyclonal MBCs clusters. **d,e,f,** Distribution of *Ighm* (**d**), *Nt5e* (CD73) and *Cd80* (**e**) and *Pdcd1lg2* (PD-L2, **f**) expression among the MSP-1 specific MBCs clusters. **g,h,** Mutation frequency of BCRs in polyclonal (**g**) and MSP-1 specific (**h**) MBCs. **i,** DEGs upregulated in clusters 0 and 2 of polyclonal MBCs and cluster 1 of MSP-1 specific MBCs.**j,k,** Gene ontology (GO) analysis of downregulated genes in MBCs from 3H3-treated mice compared to MBCs from rIgG-treated mice in clusters 0 and 2 of polyclonal MBCs combined (**j**) and cluster 1 of MSP-1 specific MBCs (**k**). Sequenced cells were pooled from 3 mice per group.

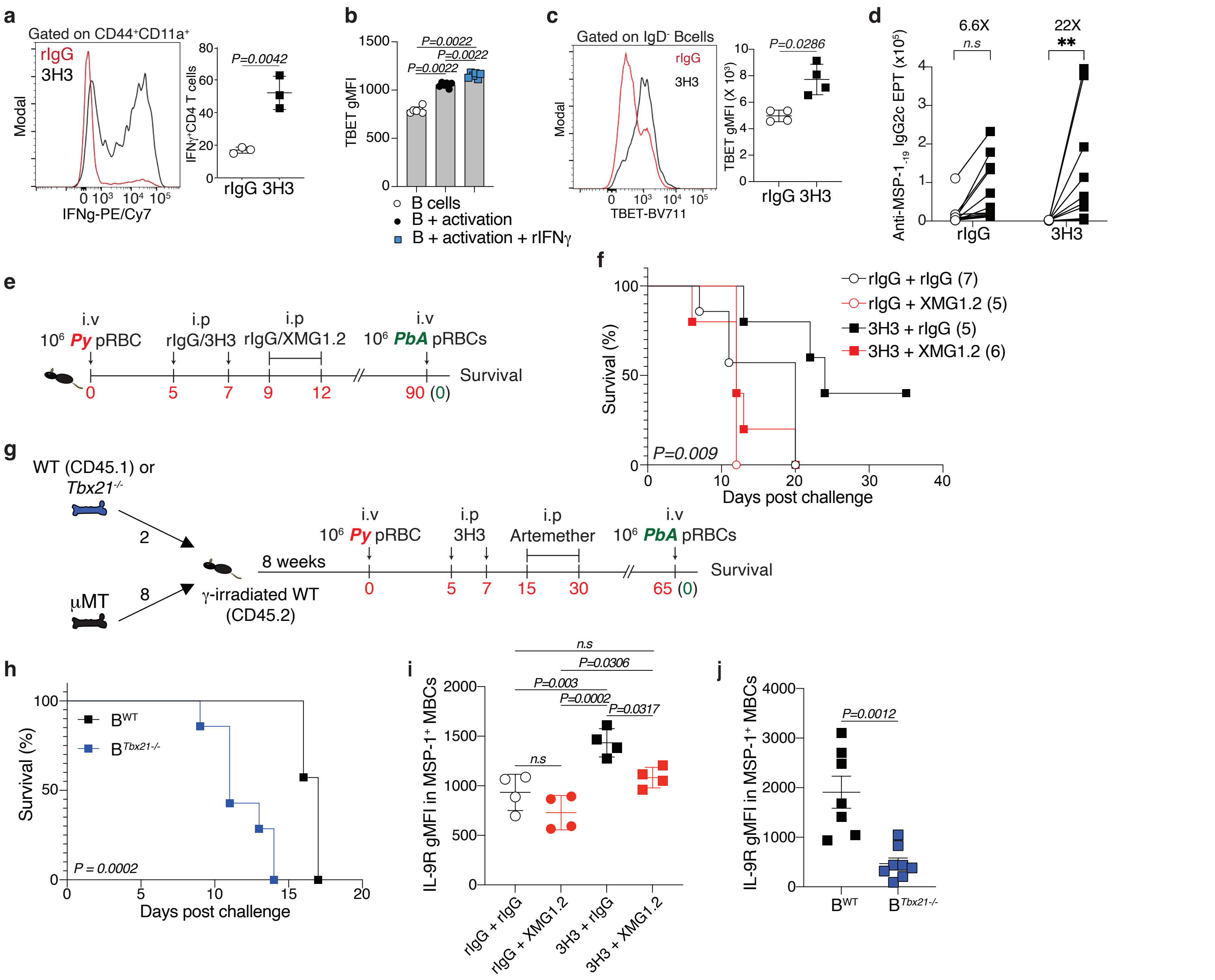


**Supp Fig. 6: 3H3-driven protection is dependent on IFNγ:T-BET:IL-9R signaling. a,** Representative histogram and frequency of CD4 T cells that produce IFNγ on 15dpi. Data are pooled from at least 2 biologically independent experiments. **b,** gMFI of TBET in B cells that were cultured for 18h in the presence or not of IFNγ. Data are pooled from 2 biologically independent experiments with 3 replicates each. **c,** Representative histogram and gMFI of TBET-expressing activated B cells on 15dpi. **d,** Fold change in anti-MSP-1-19 IgG2c serum antibody EPT between 0 and 5 dpi post *PbA* infection. Data are pooled from 2 biologically independent experiments. **e,f,** Experimental design (**e**).C57BL/6 mice were infected with *Py* and treated with 3H3 or rIgG on days 5 and 7 post infection. On days 9 to 12 post infection, mice received an IFNγ-neutralizing antibody (XMG1.2) or rIgG. During convalescence, mice were rechallenged with *PbA* and survival was monitored (**f**). Data are representative of 2 biologically independent experiments. **g,h,** Experimental design (**g**). During convalescence, mice were rechallenged with *PbA* and survival was monitored (**h**). **i,** gMFI of IL-9R in MSP-1 specific MBCs on 87-89 dpi from **e**. Data are representative of 2 biologically independent experiments and was analyzed by two-way ANOVA. **j,** gMFI of IL-9R in MSP-1 specific MBCs on 65 dpi from **g**. Data are pooled from 2 biologically independent experiments. **a**,**b**,**c**,**d** and **j** were analyzed by two-tailed Mann-Whitney U tests. **f** and **h** were analyzed by Mantel-Cox.

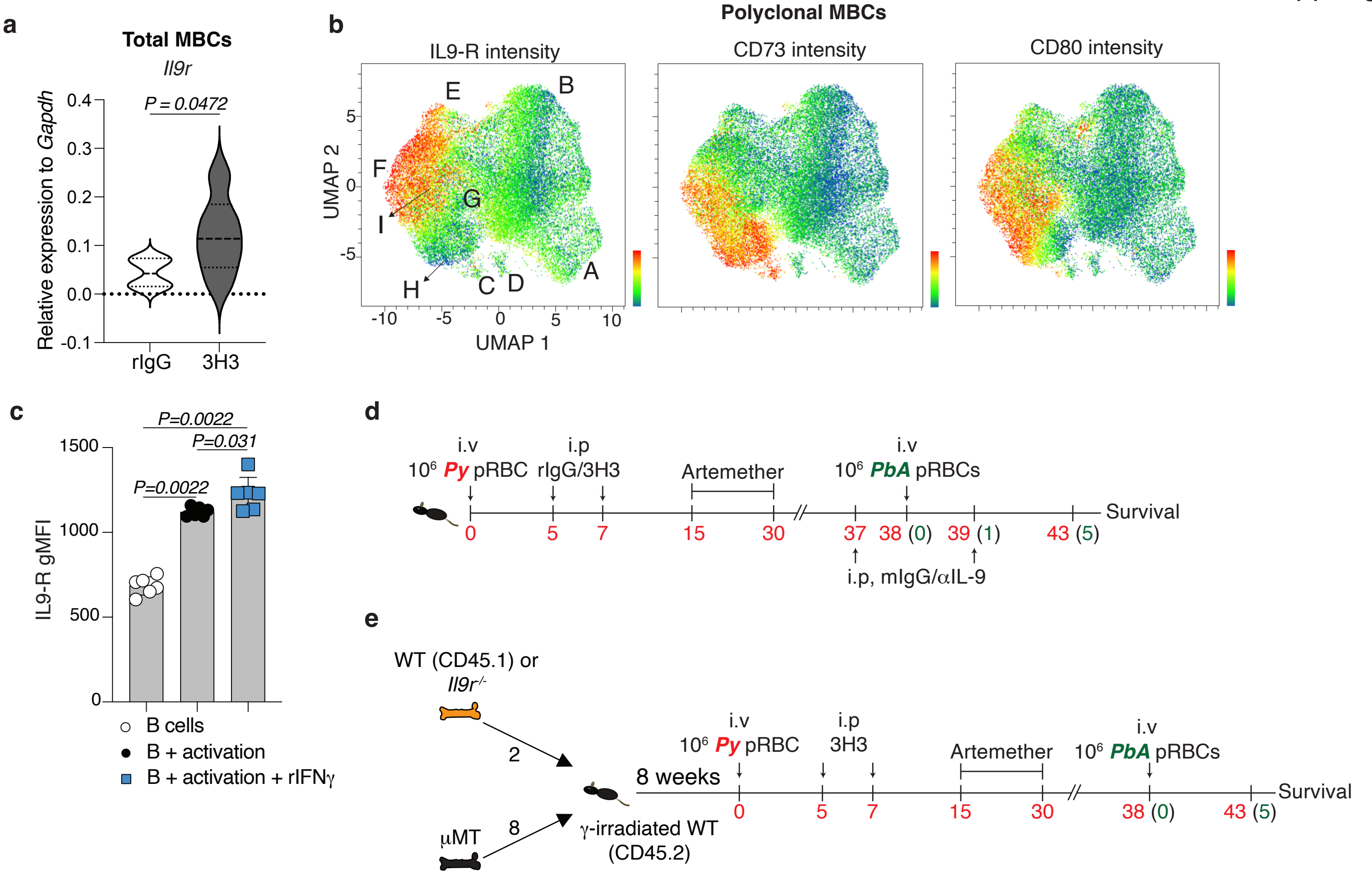


**Supp Fig. 7: 3H3-driven protection is dependent on IL-9R signaling. a,** Real-time quantitative PCR of *Il9r* normalized with *Gapdh*. Data are pooled from at least 2 biologically independent experiments. **b,** Clustering of polyclonal MBCs from FlowSOM showing IL-9R, CD73 and CD80 expression among clusters. **c,** gMFI of IL-9R in B cells that were cultured for 3 days in the presence or not of IFNγ. Data are pooled from 2 biologically independent experiments with 3 replicates each. **d,** Experimental design. C57BL/6 mice were infected with *Py* and treated with 3H3 or rIgG on days 5 and 7 post infection. One day before and after *PbA* rechallenge, mice received an IL-9-neutralizing antibody or isotype control (mIgG). During convalescence, mice were rechallenged with *PbA*. **e,** Experimental design. During convalescence, mice were rechallenged with *PbA*. **a** and **c** were analyzed by two-tailed Mann-Whitney U tests.

Supp Fig 8

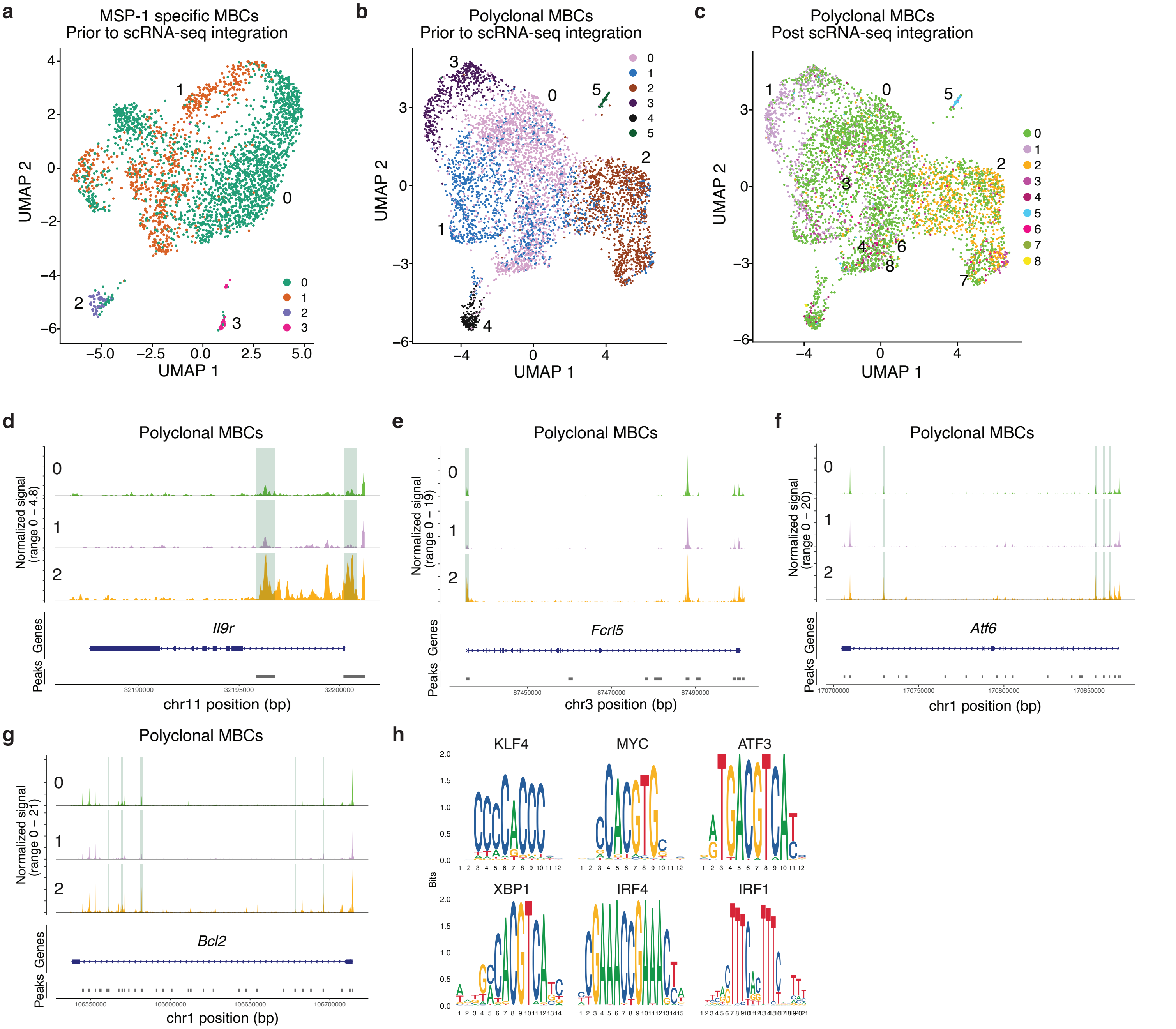


**Supp Fig. 8: Altered chromatin landscape in MBCs following 4-1BB stimulation poises them to become PCs upon recall. a,** UMAP clustering of MSP-1 specific MBCs prior to integration with scRNA-seq data. **b,c,** UMAP clustering of polyclonal MBCs prior (**b**) and post (**c**) integration with scRNA-seq data. **d-g,** *Il9r* (**d**), *Fcrl5* (**e**), *Atf6* (**f**) and *Bcl2* (**g**) peaks in clusters 0, 1 and 2 of polyclonal MBCs. **h,** Motif analysis of differentially accessible regions (DARs) in cluster 2 of polyclonal MBCs. Sequenced nuclei were pooled from 4 mice per group.
